# Defining Innate Immune Responses to the Human Gut Microbiota from Phylum to Strain

**DOI:** 10.1101/2021.11.13.468498

**Authors:** Matthew P. Spindler, Sophia S. Siu, Ilaria Mogno, Zhihua Li, Chao Yang, Saurabh Mehandru, Graham J. Britton, Jeremiah J. Faith

## Abstract

The functional potential of the gut microbiota remains largely uncharacterized. Efforts to understand how the immune system responds to commensal organisms have been hindered by the large number of strains that comprise the human gut microbiota. We develop a screening platform to measure innate immune responses towards 277 bacterial strains isolated from the human gut microbiota. We find that innate immune responses to gut derived bacteria are as strong as responses towards pathogenic bacteria, and vary from phylum to strain. Myeloid cells differentially rely upon TLR2 or TLR4 to sense particular taxa, an observation that predicts *in vivo* function. These innate immune responses can be modeled using combinations of up to 8 TLR agonists. Furthermore, the immunogenicity of strains is stable over time and following transplantation into new humans. Collectively, we demonstrate a powerful high-throughput approach to determine how commensal microorganisms shape innate immune phenotypes.

## INTRODUCTION

The gut of nearly every individual is colonized with organisms from the phyla Firmicutes, Bacteroidetes, Actinobacteria, and Proteobacteria (Qin et al., 2010). Despite the universality of these four core phyla, the gut microbiome harbors extraordinary interpersonal complexity, particular at the strain level (Eckburg et al., 2005; Faith et al., 2020, 2013; Olm et al., 2021; Schloss et al., 2014; Zhao et al., 2020). The proximity of gut bacteria to the host presents a unique challenge for the immune system, which must maintain tolerance to the resident microbiota while remaining poised to defend against infectious insults. Due in part to the immense microbial diversity within the gut microbiome, the functional relationships between specific gut microbial organisms and the host immune system are poorly understood.

Bacteria are sensed by the innate arm of the immune system through pattern recognition receptors (PRRs), which are germline encoded to sense conserved and invariant features of microorganisms (Janeway, 1989). The best characterized class of PRRs are the Toll-like receptors (TLRs), which are transmembrane receptors that recognize molecular patterns (PAMPs) contained within bacterial products. The local inflammatory response often begins when macrophages produce inflammatory cytokines, such as tumor necrosis factor (TNF)α, interleukin (IL)-1β and IL-6 in response to TLR activation (Janeway and Medzhitov, 2002). The downstream signaling pathways of PRRs are well-established, but less is known in regards to how combinations of PRRs collectively establish an optimal immune response to a particular microorganism (Ozinsky et al., 2000). Furthermore, many bacterial structures specific to particular TLR receptors remain unidentified.

Efforts to understand how bacterial species or strains within the microbiome contribute to a mucosal immune response have largely focused on the adaptive immune system (Britton et al., 2020; Britton and Faith, 2021; Geva-Zatorsky et al., 2017; Ivanov et al., 2009; Mazmanian et al., 2005; Viladomiu et al., 2017; Yang et al., 2020). Historically, the signaling pathways fundamental to our understanding of innate immunity have been elucidated using pathogen models. There are few examples of how individual members of the human gut microbiota influence innate immune responses (Seo et al., 2015; Stefan et al., 2020; Vatanen et al., 2016), and the focused scope of these examples precludes assigning general principles across taxa with regards to how gut derived bacterial strains drive innate immune profiles.

A broad understanding of the unique immunogenic features of gut derived bacterial strains is needed to better understand how the microbiome influences host immunity. Here, we utilize a scalable high-throughput implementation of an *in vitro* approach (Aleyas et al., 2009; Jang et al., 2004; Lee et al., 2014; Seo et al., 2015; Wang et al., 2010; Watkins et al., 2017) to examine innate immune cytokine responses towards a collection of 277 diverse gut bacterial strains isolated from the fecal microbiota of 17 humans. We assayed production of five cytokines in over 5,000 cocultures of gut-derived bacterial strains with mouse bone marrow derived macrophages (BMDM) or bone marrow derived dendritic cells (BMDC) (Supplementary Tables 1, 2). We found that myeloid responses to gut derived bacterial strains were highly varied, and in some cases as strong as the responses towards pathogenic bacteria. These innate immune responses varied across bacterial taxa from the phyla to the strain level. We found that responses to bacteria from the human gut microbiota could be explained using synthetic combinations of TLR agonists with both linear and non-linear synergisms, but could not be modeled by individual TLR ligands. Innate immune responses exhibited mutually-exclusive dependencies on TLR2 and TLR4 that were taxonomically defined and associated with colonic FoxP3^+^ T regulatory (Treg) cell induction in mice. We also found that *in vitro* TNFα responses towards distinct strains of *Bacteroides ovatus* correlated with fecal immunoglobulin (Ig)A production *in vivo,* collectively demonstrating that *in vitro* measurements of immunogenicity can predict specific adaptive immune responses in a colonized host. Furthermore, innate immunogenicity of human gut strains was stably maintained across human transmission and across time. Lastly, we describe a comparative genomics approach to identify bacterial genetic regions associated with strain-level variation in immunogenicity.

## RESULTS

### A Screening Platform for Measuring Innate Immune Responses to Human Gut Derived Bacteria

To gain insight into the immunogenicity of gut microbiota strains, we assembled a bacterial culture library consisting of 277 strains isolated from 8 healthy individuals, 6 individuals with Crohn’s disease (CD) and 3 individuals with ulcerative colitis (UC) (Figure 1A, Supplementary Table 3). We constructed the library to capture bacterial species that were representative of the four core phyla in the gut microbiota (Figure 1A), including multiple strains of the same species to measure variation at the closest taxonomic rank. We co-cultured BMDC and BMDM with either the filtered conditioned media of each bacterial strain grown for 48 hours or the live bacterial culture washed of metabolites, and measured secreted TNFα, IL-6, IL-10, IL-1β, and IL-23 in the coculture after 24 hours using a high-throughput ELISA workflow (Figure 1B). We selected these five analytes because of their genetic, biologic and/or therapeutic association with mucosal immunity and immune mediated diseases of the gut (Feagan et al., 2016; Hue et al., 2006; Kühn et al., 1993; Ligumsky et al., 1990; Mudter and Neurath, 2007; Targan et al., 1997; Yen et al., 2006). Our high-throughput innate screening platform was reproducible with an average R^2^ of 0.75 across biological replicates for the five analytes considered (Pearson correlation; Figure S1A). The platform also captured a broad dynamic range of dose-dependent responses to purified lipopolysaccharide (LPS, from *Escherichia coli* O55:B5), a potent TLR4 agonist (Figure S1B).

**Figure 1.**
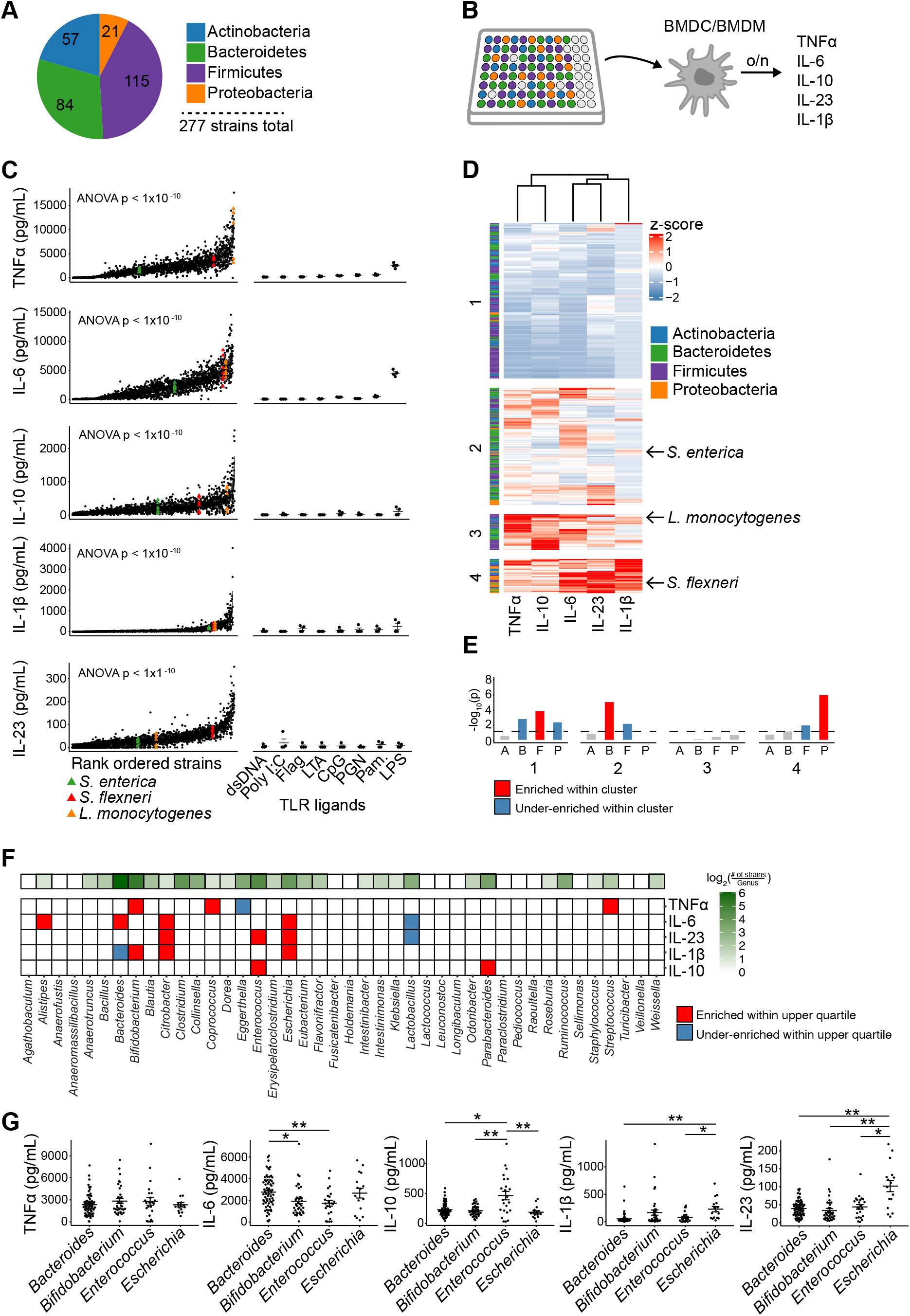
Innate Cytokine Reponses to Gut Microbial Strains Are Taxonomically Enriched and Not Explained by Single TLR Agonists. (A) A total of 277 bacterial strains spanning the four major phyla of the human gut microbiota were cultured from 8 healthy individuals, 6 individuals with Crohn’s disease (CD) and 3 individuals with Ulcerative Colitis (UC). (B) Schematic of experimental design. Strains were arrayed into plates and grown to stationary phase. A fixed volume (5 μL) of the live culture or the conditioned media was mixed with BMDC or BMDM and secreted TNFα, IL-6, IL-10, IL-1β, and IL-23 was measured after 24 hours (o/n) using a high-throughput ELISA workflow. (C) Cytokine levels as measured by ELISA from BMDC following overnight incubation with bacterial strains. Common enteric pathogens were included for reference. TLR agonist concentrations: dsDNA 10 μg/mL, poly I:C 1 μg/mL, Flag 1 μg/mL, LTA 1 μg/mL, CpG 2.5 μM, PGN 10 μg/mL, Pam 100 μg/mL, LPS 10 μg/mL. (D) Cytokine profiles of microbe-stimulated BMDC across 5 analytes. Rows were partitioned into four clusters using k-means clustering, and organized hierarchically within each cluster. Rows represent the normalized mean cytokine induction by each strain. (E) Phylum-level enrichment within the four clusters from Figure 1D. P, Proteobacteria. F, Firmicutes. B, Bacteroidetes. A, Actinobacteria. Enrichment calculated with a twosided Fisher’s exact test using membership in a phyla and membership in a cluster as the contingencies. Dotted line represents p = 0.05. (F) Enrichment of the indicated genera for higher cytokine stimulation. Enrichment was calculated with a two-sided Fisher’s exact test using membership in the genus and membership above the upper quartile as the contingencies. (G) TNFα, IL-6, IL-10, IL-1β and IL-23 responses to the four most common genera in the library. Solid horizontal lines indicate mean ± SEM. * p < 0.05, ** p < 0.01. Students t-test with Benjamini & Hochberg correction. Each point represents the mean response of an individual isolate. dsDNA, double-stranded genomic DNA from E. coli K12. Flag, flagellin. LTA, lipoteichoic acid. Pam, Pam3CSK4. PGN, peptidoglycan.

To understand innate immune responses to gut derived microbes in the context of different myeloid cells, we used a 54-member subset of the 277-strain library to compare responses to live bacterial cultures (washed with PBS and resuspended in RPMI supplemented with cysteine) between BMDM and BMDC (Supplementary table 2). We also measured responses to the conditioned media of each bacterial strain collected during stationary phase. TNFα secretion by BMDM and BMDC in response to both live cultures and conditioned media were significantly correlated (live cultures R^2^ = 0.28, p = 9.30×10^-6^, conditioned media R^2^ = 0.69, p < 1×10^-10^, Pearson correlation; Figures S1C, S1D), with a proportionally higher signal observed in BMDC. TNFα secretion in response to live bacterial cultures did not correlate with responses to conditioned media (Figures S1E, S1F). Importantly, the assay was robust to variation in bacterial growth density across strains in the library, as TNFα secretion only weakly correlated with the optical density (600nm) of the bacterial culture (R^2^ = 0.02, p = 0.02, Pearson correlation; Figure S1G). Based on these initial metrics, we focused our analysis on BMDC responses towards live cultures of gut derived bacteria since this model produced a larger dynamic range across all analytes considered, was more sensitive and more reproducible, and elicited a robust IL-23 response (Supplementary Table 2).

### Innate Cytokine Reponses to Gut Microbial Strains Are Taxonomically Enriched and Not Explained by Single TLR Agonists

BMDC responses towards live cultures of gut derived bacteria led to a broad range of cytokine profiles that significantly varied by several orders of magnitude across strains (TNFα ANOVA p < 1×10^-10^; Figure 1C) by several orders of magnitude. To place these responses in the context of known stimuli, we considered how innate cytokine responses to gut derived bacteria compared to responses towards 8 common purified TLR ligands. We calculated dose-response relationships with each of these individual TLR agonists, and selected concentrations which elicited the highest response or at which the signal saturated (Figure S1H). Stimulating BMDC with these common TLR ligands did not capture the range of responses observed upon stimulation with the commensal bacterial library (Figure 1C). We also compared gut-derived bacterial responses to three well-characterized enteric pathogens *(Listeria monocytogenes, Salmonella enterica* and *Shigella flexneri*). The cytokine responses to these pathogens largely fell in the upper part of the cytokine distributions (Figure 1C), suggesting that innate responses towards the majority of commensal isolates are on average lower than responses towards common enteric pathogens. However, all cytokine responses to these common pathogens were contained within the range observed across the 277-strain library (Figure 1C). Together, these results indicate that innate responses to gut derived bacterial strains can be as strong as responses towards pathogenic bacteria, span a tremendous dynamic range, and cannot be reproduced by individual TLR ligands.

Given long-standing efforts to identify commensal strains that contribute to inflammatory responses in inflammatory bowel diseases (IBD) (Onderdonk et al., 1987; Rath et al., 1999, 1996; Seo et al., 2015), we tested if commensal strains isolated from patients with UC and CD elicited specific cytokine signatures. We found no enrichment for greater cytokine responses in IBD-associated bacterial strains compared to strains derived from healthy donors (Figure S1I), and there were no significant differences in the overall mean level of TNFα, IL-6, IL-10, IL-1β and IL-23 produced in response to healthy donor strains versus strains derived from UC or CD donors (Figure S1J). Broadly, these data suggest that specific innate immune responses towards individual gut derived bacterial strains are not enriched by IBD status.

To examine if certain bacterial taxa associate with the induction of particular cytokine signatures, we applied common clustering and dimensional reduction algorithms. K-means clustering of cytokine responses (k = 4; Figure S1K) segregated bacteria into four groups that broadly associated with phyla (Figure 1D). Firmicutes were enriched in a cluster encompassing the lowest cytokine responses across all analytes (Cluster 1 Firmicutes p = 1.51×10^-4^, Fisher’s exact test; Figure 1E), while Bacteroidetes were enriched in a cluster comprising a milder cytokine phenotype (Cluster 2 Bacteroidetes p = 8.92×10^-6^, Fisher’s exact test; Figure 1E). Proteobacteria were enriched in a cluster with the highest IL-1β and IL-23 induction (Cluster 4 p = 1.24 x10^-6^, Fisher’s exact test; Figures 1E), and promoted greater secretion of IL-1β and IL-23 compared to the other phyla in the library (p < 0.01, Student’s t-test with Benjamini & Hochberg (BH) correction; Figure S1L). Principal component analysis discriminated between a colinear TNFα/IL-10 axis and an orthogonal IL-23/IL-1β Proteobacteria driven response (Figure S1M).

We next looked for genera within the library that were enriched in the top extremes of the cytokine responses (using the 75^th^ quantile as a cutoff; Figure 1F). Focusing on the four genera with the highest number of strains per genus in the library (*Bacteroides* n = 67 strains, *Bifidobacterium* n = 37 strains, *Enterococcus* n = 23 strains and *Escherichia* n = 15 strains), we observed that *Enterococcus* members were enriched for IL-10 induction (Figure 1F), and induced on average more IL-10 than *Bacteroides, Bifidobacterium* and *Escherichia* (p < 0.05, p < 0.01, and p < 0.01 respectively, Student’s t-test with BH correction; Figure 1G). Members of *Escherichia* were enriched for strains in the upper quartile of IL-6, IL-23, and IL-1β induction (Figure 1F), and induced more IL-23 than *Bacteroides, Bifidobacterium* and *Enterococcus* (p < 0.01, p < 0.01, and p < 0.05 respectively, Student’s t-test with BH correction; Figure 1G), which is consistent with studies suggesting these bacteria can elicit Th17-biased immunity *in vivo* (Atarashi et al., 2015; Britton et al., 2020; Viladomiu et al., 2017). *Bacteroides* were enriched within strains that elicited high levels of IL-6 (Figure 1F), and IL-6 production in response to *Bacteroides* was higher than towards *Bifidobacterium* and *Enterococcus* (p < 0.05 and p < 0.01 respectively, Student’s t-test with BH correction; Figure 1G). Together, these results demonstrate that innate immune responses towards gut microbial strains vary across taxa, with broad enrichments in cytokine response at the phylum level and more refined enrichments at the genus level.

### Combinations of TLR Ligands Reflect Cytokine Responses Towards Gut-Derived Bacterial Stains

Although macrophages and dendritic cells express TLRs (Reis E Sousa, 2004; Visintin et al., 2001), it is unclear how the cell responds to the complex mixture of PAMPs on the surface of bacteria. To more accurately model a scenario in which a cell responds to a bacterial organism, we assembled from a set of eight TLR agonists all 255 possible combinations of 8 choose *K*(8 choose 1, 8 choose 2, etc…; Figures 2A, 2B). Responses following stimulation by these 255 different TLR ligand combinations increased on average with the value of *K* (TNFα ρ = 0.55, ρ < 1×10^-10^, IL-6 ρ = 0.52, p < 1×10^-10^, IL-10 p = 0.51, p < 1×10^-10^, IL-1β ρ = 0.34, p = 2.48×10^-8^, Spearman correlation; Figures 2A, 2C). We found that combinations of three TLR agonists (*K* = 3) elicited the largest range of responses (Figure S2A), and that as *K* increased the variance in responses decreased (Figure S2B). Overall, the response to these 255 different TLR ligand combinations could explain the vast majority of the BMDC responses to the 277 bacterial strains across all analytes (Figures 2D, S2C), providing an opportunity to use these synthetic TLR combinations to gain insights into how bacteria elicit particular cytokine responses.

**Figure 2.**
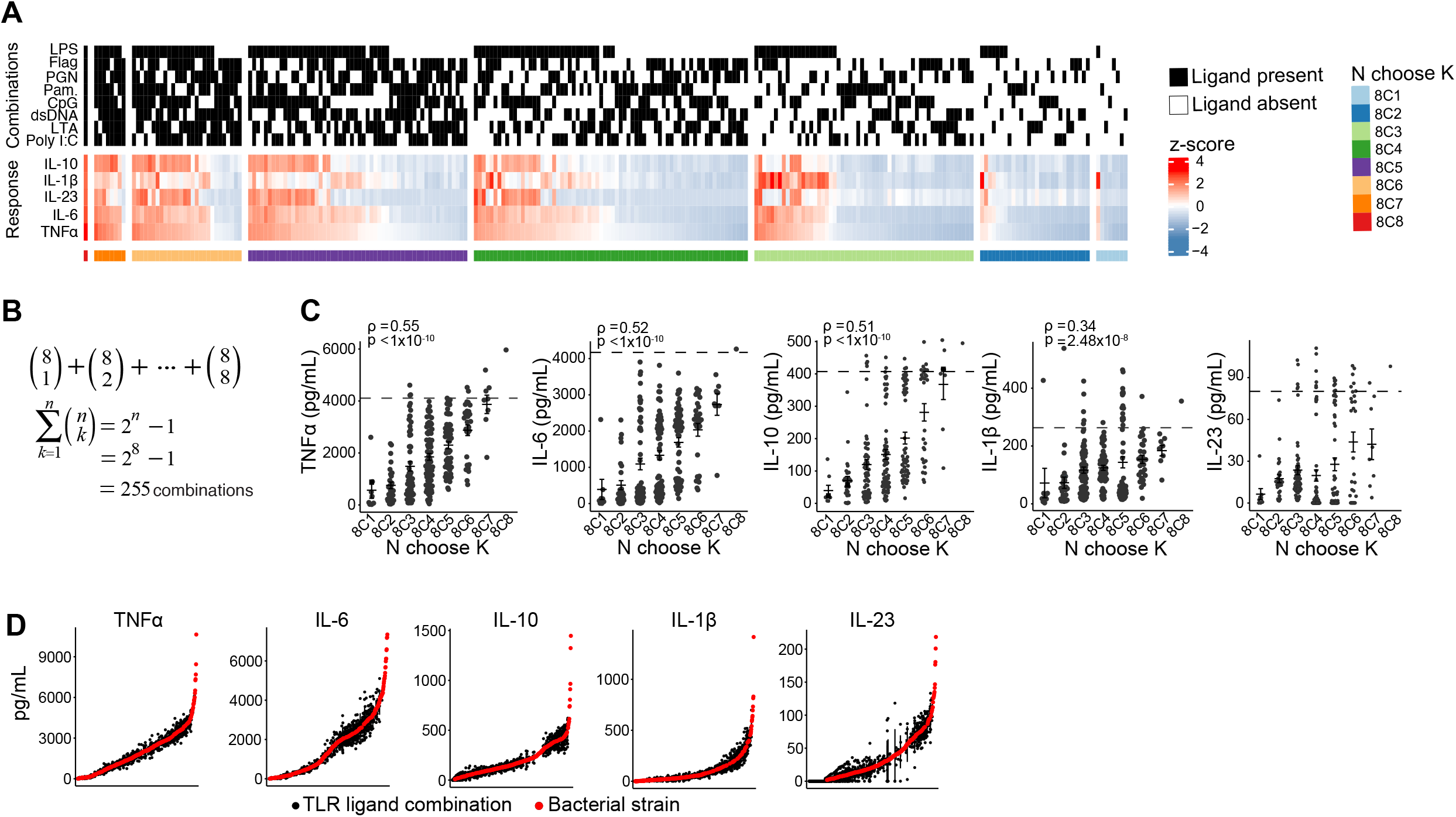
Combinations of TLR Ligands Reflect Cytokine Responses Towards Gut-Derived Bacterial Stains. (A) Heatmaps relating the presence or absence of a TLR ligand in every combination assayed (255 in total) with the corresponding normalized mean cytokine value elicited from BMDC. (B) Schematic of experimental design. Eight artificial TLR ligands were assembled in every possible combination of 8 (a total of 255 combinations). (C) Cytokine responses increase as a function of *K* (combination complexity). Dotted lines represent the 90^th^ quantile in the distribution of responses to a commensal bacterial collection (Figure 1C). Correlation coefficient determined by Spearman’s rank correlation. (D) Responses to 255 unique TLR ligand combinations explain responses to 277 unique gut derived bacteria. dsDNA, double-stranded genomic DNA from E. coli K12. Flag, flagellin. LTA, lipoteichoic acid. Pam, Pam3CSK4. PGN, peptidoglycan.

### Pairwise Synergies Between TLR Ligands Boost Cytokine Responses

We first assessed the individual impact of each TLR ligand on cytokine induction using statistical modelling techniques. Unsurprisingly, LPS had the largest effect (TNFα p < 0.0001, Student’s t-test), followed by cytosine guanosine dinucleotide (CpG) (TNFα p < 0.0001, Student’s t-test) and Pam3CSK4 (TNFα p < 0.0001, Student’s t-test; Figures 3A, S3A). To uncover synergy between TLR agonists and cytokine response, we used stepwise linear regression and elastic net regression to model pairwise interactions in our dataset. Both stepwise and elastic net regressions generated similar coefficient estimates (R^2^ = 0.86, p = 1×10^-10^, Pearson correlation; Figure S3B). We found synergy between LPS and flagellin (Figure 3B, blue box) in the production of TNFα (p = 0.019, Student’s t-test with BH correction), consistent with studies suggesting that TLR4 and TLR5 physically associate to trigger inflammatory responses (Hussain et al., 2020). LPS and CpG also acted synergistically (Figure 3B, red boxes) to enhance the production of TNFα, IL-6, IL-10 and IL-23 (TNFα, IL-6, IL-10, IL-23 *p* < 0.0001, Student’s t-test with BH correction, Figure 3C), which corroborates previous work demonstrating that TLR4 and TLR9 non-additively amplify cytokine signaling (De Nardo et al., 2009; Napolitani et al., 2005). Notably, LPS and CpG together reduced the level of IL-1β compared to LPS alone (*p* < 0.001, Student’s t-test with BH correction; Figure 3C), perhaps via an IL-10 negative feedback mechanism (Abrams et al., 1991).

**Figure 3.**
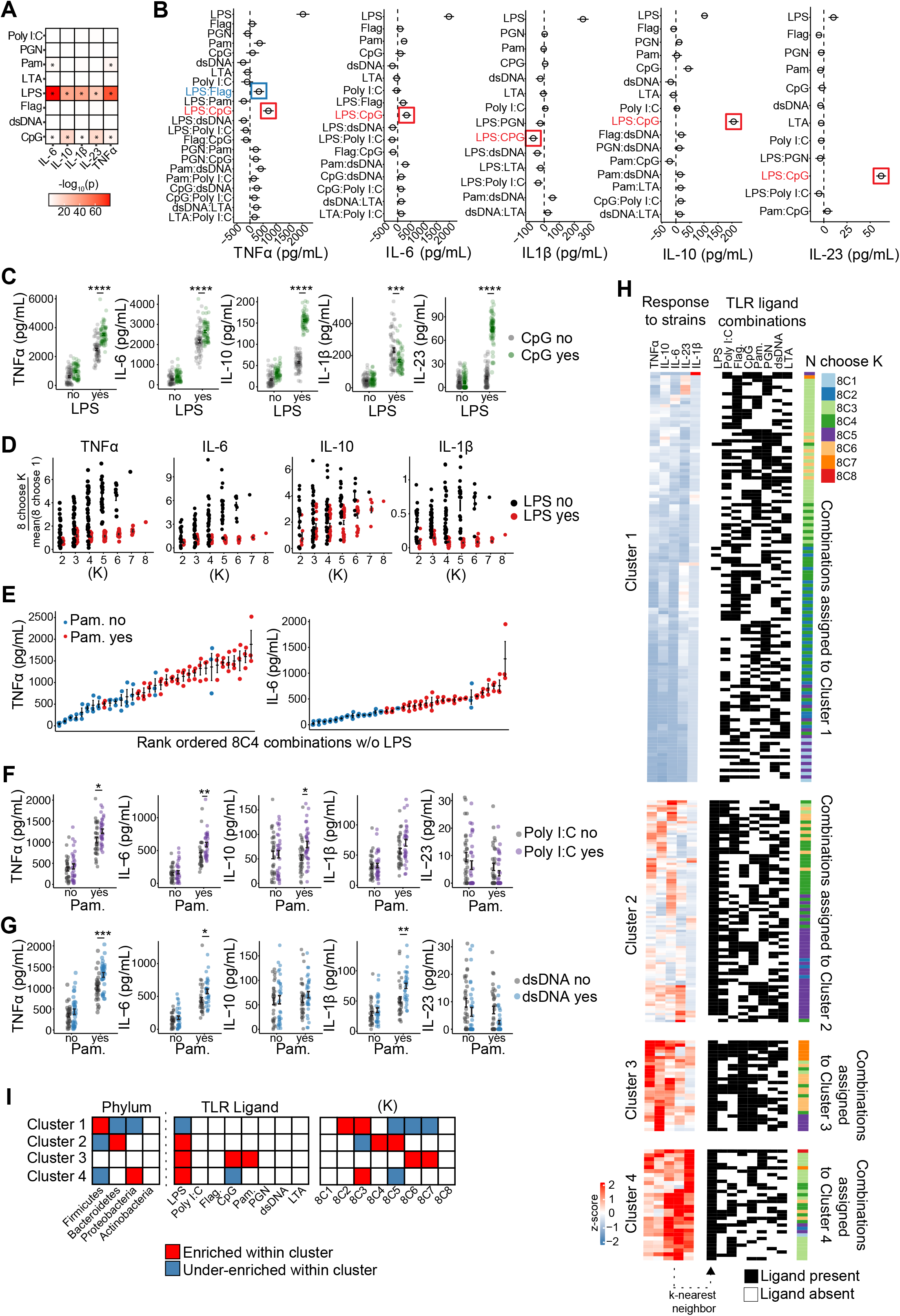
Pairwise Synergies Between TLR Ligands Boost Cytokine Responses. (A) Comparisons of cytokine stimulation in the presence or absence of the indicated TLR ligands (Student’s t-test). * represents p < 0.05. (B) Summary of stepwise linear regression used to identify pairwise interactions between TLR ligands that synergistically enhance cytokine secretion in BMDC. (C) TNFα production in the presence or absence of LPS and CpG. *** p < 0.001, **** p < 0.0001. Student’s t-test with BH correction. (D) Fold change as a function of *K* in the presence or absence of LPS. Fold change calculated by normalizing to the mean of N choose 1. (E) Rank ordered TNFα and IL-6 responses to TLR ligand combinations of 8 choose 4 with conditions containing LPS removed. (F) Responses to Pam3CSK4 in the presence or absence of poly I:C. ** p < 0.01, * p < 0.05. Student’s t-test with BH correction. (G) Responses to Pam3CSK4 in the presence or absence of dsDNA. *** p < 0.001, ** p < 0.01, * p < 0.05. Student’s t-test with BH correction. (H) Cytokine responses to 277 unique bacterial strains were clustered using k-means clustering (Figure 1D), and used as a training dataset to classify responses to 255 different TLR ligand combinations using the k-nearest neighbor algorithm. (I) Left side: Phylum enrichment (Fisher’s exact test) in indicated cluster (Figure 1E). Right side: Enrichment (Fisher’s exact test) in the indicated cluster by TLR ligand or by *K* (combination complexity). dsDNA, double-stranded genomic DNA from E. coli K12. Flag, flagellin. LTA, lipoteichoic acid. Pam, Pam3CSK4. PGN, peptidoglycan.

To uncover other nonadditive effects between TLR ligands, we segregated our data into conditions where LPS was present or absent. There was a larger dynamic range in response to ligand combinations which did not include LPS (Figure 3D), suggesting that 1) the presence of LPS drives a saturation of the cytokine response, and 2) innate immune responses can be exquisitely scaled depending on which ligands are present. Pam3CSK4 had a large individual effect when LPS was excluded (Figures 3E, S3C), implicating TLR2 as an important signaling potentiator in our model system. We found significant synergy between Pam3CSK4 and poly I:C (a TLR3 ligand) in the induction of TNFα, IL-6 and IL-10 (TNFα p < 0.05, IL-6 p < 0.01, IL-10 p < 0.05, Student’s t-test with BH correction; Figure 3F). TLR2 and TLR9 crosstalk is thought to contribute to host defense against pathogenic bacteria and parasites (Bafica et al., 2006, 2005), and we found that double-stranded DNA from *E. coli* (a TLR9 ligand) and Pam3CSK4 together enhanced TNFα, IL-6, and IL-1β (TNFα p < 0.001, IL-6 p < 0.05, IL-1β p < 0.01, Student’s t-test with BH correction; Figure 3G). Notably, CpG (also a TLR9 ligand) and Pam3CSK4 did not synergize, suggesting that ligand specific factors may be important in regulating the scale of the response (Iwasaki and Medzhitov, 2010; Long et al., 2009). Together, these data suggest that pairwise synergies between TLRs occur in the context of complex combinations of TLR agonists.

To gain insight into the types of TLR ligand combinations that correspond to particular groups of gut derived bacteria, we used supervised machine learning to classify the response of each TLR ligand combination using responses to the 277-member bacterial strain library as a training dataset. Specifically, we assigned each TLR ligand combination a membership to one of the four clusters that broadly associated with each phylum (Figure 1C) using a k-nearest neighbor algorithm (Figure 3H). We found that ligand combinations containing LPS were predominantly assigned into clusters 2, 3 and 4, which associated with responses to Bacteroidetes and Proteobacteria (Figures 3H, 3I). The complexity (*K*) of the TLR ligand combination also factored into assignments, as combinations of *K* = 2 and *K* = 3 were enriched in the Firmicutes associated cluster (i.e., cluster 1), while combinations of *K* = 4 and *K* = 5 were enriched in the Bacteroidetes associated cluster (i.e., cluster 2; Figure 3I). These analyses suggest that the absence of LPS with low-complexity TLR ligand combinations best replicate a response to Firmicutes, while the presence of LPS plus intermediate-complexity TLR ligand combinations broadly explain responses to Bacteroides. More generally, TLR ligand combinations can be engineered to replicate responses to specific bacterial taxa, suggesting that defined TLR stimulation can model aspects of innate immune responses to bacteria from the human gut microbiota.

### Sensing of Gut Derived Bacteria by TLR2 and TLR4 is Taxonomically Defined

To determine the immune receptors necessary for the innate immunogenicity of gut derived bacteria, we subsampled the 277-member commensal library into a smaller 54-member library containing representatives from all four main gut phyla and all four functional clusters (Figure 1), and screened responses using BMDC derived from TLR knockout (KO) animals. First, we confirmed that wild-type BMDC responses to the subsampled 54-member library correlated with responses to the original 277-member library (Figure S4A). We found the production of cytokines in response to gut derived bacteria was completely abrogated in BMDC deficient in MyD88, the signaling adaptor protein used by many cell surface TLRs, including TLR2 and TLR4 (p < 0.0001, paired Student’s t-test; Figure S4B; Hou et al., 2008). Provided that TLR2 and TLR4 ligands were also strong potentiators of cytokine induction in the combinatorial TLR ligand system (Figures 2, 3), we explored innate immune responses to gut microbes using BMDC deficient in these respective genes (TLR2^KO^ or TLR4^KO^; Wooten et al., 2002). To validate our KO system, we measured the dose-response to LPS and confirmed that cytokine levels diminished (Figure S4C). We then examined responses to gut derived bacteria using TLR2^KO^ and TLR4^KO^ BMDC and compared these to responses by wildtype BMDC. Using hierarchical clustering, we grouped the fold-change per bacterial strain over all cytokines and over all KOs, and found that strains segregated by phyla (Figure 4A). Actinobacteria and Firmicutes were largely resistant to the loss of TLR2 or TLR4 (Figures 4B, S4D), while responses to Proteobacteria depended on TLR4 but not TLR2 (p < 0.001, Students t-test; Figures 4B, S4D). In contrast, IL-6 and IL-10 secretion in response to Bacteroidetes decreased with the loss of either TLR2 or TLR4 (IL-6 TLR2 p < 0.001, IL-6 TLR4 p < 0.001, Students t-test; Figures 4B, S4D).

**Figure 4.**
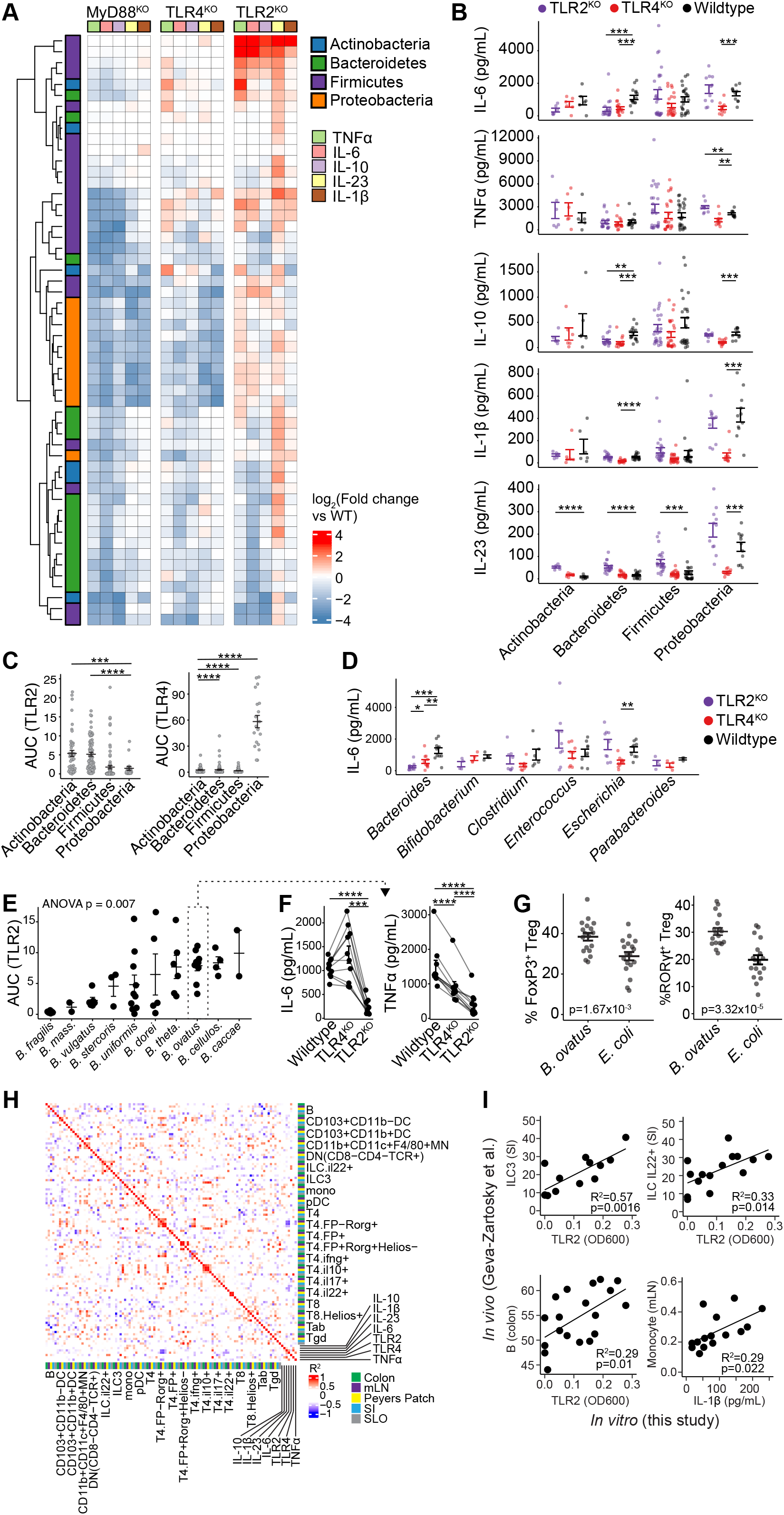
Sensing of Gut Derived Bacteria by TLR2 and TLR4 is Taxonomically Defined. (A) Fold-change of knock-out responses compared to wildtype. Each row represents a unique isolate. Rows are clustered hierarchically. A threshold was applied to very low measurements as described in the methods. (B) Cytokine responses to indicated phylum in BMDC. ** p < 0.01, *** p < 0.001, **** p < 0.0001. Student’s t-test with BH correction. (C) Bacterial strains from distinct phyla differentially stimulate TLR2 and TLR4. TLR stimulation measured by HEK-Blue murine TLR reporter cell lines. *** p < 0.001, **** p < 0.0001. Student’s t-test with BH correction. Each point represents the response of an individual isolate. (D) IL-6 secretion to bacteria from the indicated genera in wildtype, TLR2^KO^ and TLR4^KO^ BMDC. Each point represents the mean response to a particular isolate. * p< 0.05, ** p < 0.01, *** p < 0.001. Student’s t-test with BH correction. (E) TLR2 stimulation varies across *Bacteroides* species. Stimulation measured by HEK-Blue murine TLR reporter cell lines. Each point represents the response to an individual isolate. (F) IL-6 and TNFα production in response to *B. ovatus* in wildtype, TLR2^KO^ and TLR4^KO^ BMDC. Lines connect identical strains. *** p < 0.001, **** p < 0.0001. Paired student’s t-test with BH correction. (G) The proportion of FoxP3^+^ Treg cells and RORgt+ FoxP3+ Treg cells in the colon of mice monocolonized with strains of *B. ovatus* or strains of *E. coli.* Each point represents one mouse. 6 strains per species, and 3 mice per strain were used. Student’s t-test. (H) Integration of data from a previously published dataset (Geva-Zatorsky et al., 2017). Correlation matrix between immune cell populations in the indicated tissues, cytokine secretion, and TLR stimulation across 21 species present in both studies. Multiple strains of the same species were averaged. Pearson correlation. (SI, small intestine. mLN, mesenteric lymph node. SLO, secondary lymphoid organ). (I) Comparisons between *in vivo* immune populations and *in vitro* measurements across 21 species present in both studies (Figure 4H). Pearson correlation. Regression p-values calculated by F test.

To gain further insight into the engagement of specific TLRs by gut derived bacteria, we assembled conditioned media from members of the full 277-member bacterial strain library, and assayed TLR2 and TLR4 stimulation using HEK-Blue reporter cell lines expressing murine TLR2 or TLR4. We found that Actinobacteria and Bacteroidetes broadly stimulated TLR2 (Figure 4C), while Proteobacteria selectively stimulated TLR4 (p < 0.0001, Student’s t-test with BH correction; Figure 4C). Few bacteria in the collection stimulated both TLR2 and TLR4 (R^2^ = 0.01, p = 0.13, Pearson correlation; Figure S4E).

We next compared responses to the genera most represented in the bacterial library using TLR2^KO^ and TLR4^KO^ BMDC. *Clostridium* and *Enterococcus* did not depend on either TLR2 or TLR4 (Figure 4D), consistent with the observation that members of these two genera did not stimulate TLR2 or TLR4 (Figures 4C, S4F). Interestingly, *Bifidobacterium* stimulated TLR2 (Figure S4G), but IL-6 secretion in response to *Bifidobacterium* was not significantly affected by the absence of TLR2, suggesting redundancy in the sensing of this genus (Figure 4D). In contrast, IL-6 and IL-10 responses to *Bacteroides* were strongly TLR2 dependent (IL-6 p < 0.001, IL-10 p < 0.01, Student’s t-test with BH correction; Figures 4D, S4H), and IL-6 responses to *Bacteroides* were more dependent on TLR2 than on TLR4 (IL-6 p < 0.05, Student’s t-test with BH correction; Figure 4D). Meanwhile, responses to *Escherichia* were lost in TLR4-deficient cells (IL-6 p < 0.01, TNFα p < .05, IL-10 p < 0.01, Student’s t-test with BH correction), but were unaffected by TLR2 deficiency (Figures 4D, S4H).

We next looked at the TLR2 stimulation potential within species of *Bacteroides.* We found that *Bacteroides* species exhibited a range of TLR2 stimulation (p = 0.007, ANOVA), with *B. ovatus* among those which most highly stimulated TLR2 (Figure 4E). Using a collection of 10 distinct *B. ovatus* strains, we found that IL-6 secretion in response to *B. ovatus* strains was indeed TLR2 dependent (p < 0.001, paired Student’s t-test with BH correction), and not dependent on TLR4 (Figure 4F). Furthermore, the loss of TLR2 diminished TNFα secretion in response to *B. ovatus* strains more than the loss of TLR4 (p < 0.0001, paired Student’s t-test with BH correction; Figure 4F).

Since *B. ovatus* is sensed by TLR2, we reasoned that *B. ovatus* may drive TLR2-mediated immunity *in vivo.* Blockade of TLR2 *in vivo* diminishes the number of colonic IL-10 producing lymphocytes (Mishima et al., 2019). Furthermore, the induction of FoxP3^+^ Treg cells is diminished in TLR2^KO^ mice (Netea et al., 2004) and augmented by the application of TLR2 agonists (Round and Mazmanian, 2010). Therefore, we hypothesized that *B. ovatus* would be a potent inducer of intestinal FoxP3^+^ Treg cells. We compared the levels of FoxP3^+^ Treg cells in the colonic lamina propria of mice monocolonized with strains of *E. coli,* which is sensed by TLR4 and not TLR2 (Figure S4I), to mice monocolonized with strains of *B. ovatus.* We found that all *B. ovatus* strains potently induced FoxP3^+^ Treg cells in the colon, and that the levels of FoxP3^+^ Treg cells were significantly higher in mice monocolonized with *B. ovatus* than in mice monocolonized with *E. coli* (p = 1.67×10^-3^, Student’s t-test; Figure 4G). Furthermore, the levels of colonic RORγt^+^ Treg cells were also higher in mice monocolonized with *B. ovatus* compared to mice monocolonized with *E. coli* (p = 3.32×10^-5^, Student’s t-test; Figure 4G). Taken together, these data suggest there are phylum-level distinctions in innate immune sensing of commensal microbes by TLR2 and TLR4. Within phyla that stimulate TLR2, there exists broad variation that can be explained by genus and species level enrichments. Furthermore, species level differences in TLR2 and TLR4 recognition associate with alterations in colonic FoxP3^+^ Treg cell induction.

To further correlate our *in vitro* observations with the *in vivo* functionality of the human gut microbiota, we integrated data from an existing study that systematically analyzed host immune phenotypes following monocolonization of mice with diverse human gut-derived bacteria (Geva-Zatorsky et al., 2017). We correlated *in vivo* immunologic phenotypes in the gut and secondary lymphoid organs with our *in vitro* cytokine and TLR stimulation measurements across 21 species included in both datasets (Figure 5H). We found several significant associations between our *in vitro* data and *in vivo* immune phenotypes driven by species variation within the microbiota. These included associations between *in vitro* TLR2 stimulation and the frequency of colonic B cells (R^2^ = 0.29, p = .01, Pearson correlation), TLR2 stimulation and the frequency of both innate lymphoid cell (ILC)3 and IL-22^+^ ILC in the small intestine (R^2^= 0.57, p = 0.0016 and R^2^= 0.33, p = 0.014, Pearson correlation), and between *in vitro* induced IL-1β and the frequency of monocytes in the mesenteric lymph node (mLN) (R^2^= 0.29, p = 0.022, Pearson correlation; Figure 4I). These analyses demonstrate the power of cross-referencing large high-dimensional datasets to better characterize the function of the human gut microbiota, and suggest that high-throughput *in vitro* screening approaches can predict *in vivo* immune phenotypes across species of the human gut microbiota.

**Figure 5.**
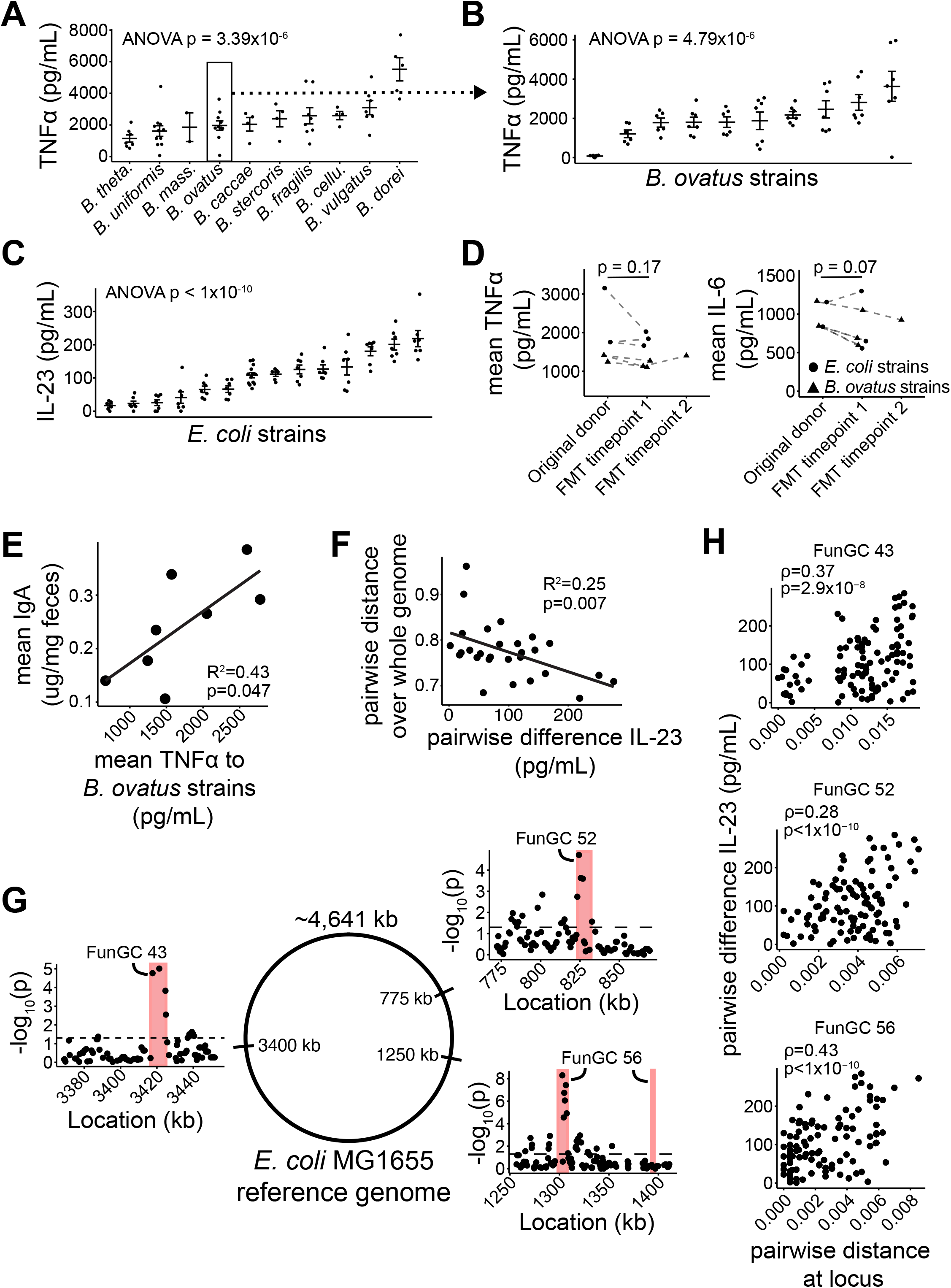
Differences between gut derived bacterial strains drive variation in innate immunogenicity. (A) TNFα induction varies across Bacteroides species. Each point represents the mean response of an individual isolate. (B-C) Cytokine responses across strains of *B. ovatus* and *E. coli.* Each point represents a replicate condition. (D) TNFα and IL-6 induction is stably maintained across human transmission and across time. Dotted lines connect the original donor with an FMT recipient. Two donors had multiple recipients. One recipient was resampled at two timepoints. FMT timepoint 1 corresponds to 2-3 months following transplant, FMT timepoint 2 corresponds to 11 months following transplant. (E) TNFα stimulation in response to *B. ovatus* strains correlates with fecal IgA measurements from gnotobiotic mice monocolonized with the corresponding *B. ovatus* strains. Mean TNFα measurements represent the average of four biologic replicates. Mean IgA measurements represent an average from 4-8 mice per strain. Regression p values calculated by F-test. (F) Pairwise distances over the *E. coli* genome negatively correlates with the pairwise difference in IL-23 induction within closely related strains of *E. coli* (greater than 65% sequence homology, Figure S5E). (G) Strong associations between protein distances within consecutive 5-gene windows and IL-23 induction coalesce at genomic locations that correspond with functional genomic clusters (FunGCs). Red rectangles represent the genomic borders of the indicated FunGC. (H) Correlations between the pairwise protein distance across consecutive genes and the pairwise difference in the capacity to induce IL-23 at the genomic locations corresponding to the indicated FunGC. Spearman rank correlation.

### Differences between gut derived bacterial strains drive variation in innate immunogenicity

Having identified broad taxa that associate with innate cytokine responses and species-level enrichments in TLR2 potentiation, we next assessed functional variation between more closely related isolates. We focused our analyses on the gram negative genera *Bacteroides* and *Escherichia* given their near ubiquity in the human gut microbiome, our data suggesting a differential reliance on TLR2 and TLR4 signaling, and previous studies showing non-uniform effects of *B. ovatus* and *E. coli* strains on the mucosal adaptive immune system (Atarashi et al., 2015; Britton et al., 2020; Viladomiu et al., 2017; Yang et al., 2020). First, we measured cytokine induction by multiple strains of *Bacteroides* across 10 different species. We observed significant variation in TNFα, IL-6, and IL-10 induced by different species of *Bacteroides* (TNFα p = 3.39×10^-6^, IL-6 p = 0.004, IL-10 p = 0.016, ANOVA; Figures 5A, S5A). When comparing cytokine induction across 10 distinct strains of *B. ovatus,* the induction of TNFα, IL-6, and IL-10 also varied significantly (TNFα p = 4.79×10^-6^, IL-6 p = 1.98×10^-10^, IL-10 p = 8.33×10^-5^, ANOVA; Figures 5B, S5B). In addition, the induction of TNFα, IL-23, IL-6, IL-10 and IL-1β varied significantly across 14 distinct strains of *E. coli* (TNFα, IL-23, IL-6, and IL-10 p < 1×10^-10^, IL-1β p = 2.33×10^-10^, ANOVA; Figures 5C, S5C). Together, these data demonstrate that different strains of the same species of bacteria derived from the human gut microbiota drive varied innate immune responses.

Several *E. coli* and *B. ovatus* strains present in our library originated from stool that was later transferred to human recipients with recurrent *Clostridioides difficile* infection as fecal microbiota transplantation (FMT) therapy (Aggarwala et al., 2021). We tested if strain immunogenicity was maintained following transplantation into a recipient human host by comparing responses to strains isolated from the donor microbiota before transplant with responses to the same strain re-isolated from the recipient microbiota post-transplant. We found no difference in the mean TNFα and mean IL-6 induced by strains isolated from the original donor compared to strains isolated from the recipient following FMT (TNFα p = 0.17, IL-6 p = 0.07, paired Student’s t test; Figure 5D). Of the 6 donor-recipient pairs examined, 5 out of 6 elicited similar TNFα induction, and 6 out of 6 elicited similar IL-6 following re-isolation in the recipient 2-3 months post-FMT (FMT timepoint 1; Figure 5D). One FMT recipient was sampled longitudinally, and the same *B. ovatus* strain re-isolated 11 months post-FMT elicited comparable TNFα and IL-6 induction to the isolate from the original donor (FMT timepoint 2; Figure 5D), suggesting that innate immunogenicity of human gut strains is stably maintained across human transmission and across time.

We considered whether differences between gut derived bacterial strains associate with *in vivo* immune phenotypes. *B. ovatus* is a known inducer of fecal IgA, and for reasons unknown, some strains of *B. ovatus* are capable of inducing higher IgA than other *B. ovatus* strains (Yang et al., 2020). We found that strain-level *in vitro* TNFα responses to *B. ovatus* correlated with *in vivo* fecal IgA levels in gnotobiotic mice monocolonized with the corresponding *B. ovatus* strain (R^2^ = 0.43, p = 0.047, Pearson correlation; Figure 5E). Of note, we did not find a significant correlation between IL-6 in response to *B. ovatus* and fecal IgA levels (Figure S5D). This suggests that strain level differences measurable *in vitro* may translate to functional and predictable effects on immune responses *in vivo.*

One of the most striking examples of inter-strain variation that we identify is the varied induction of IL-23 across strains of *E. coli* (Figure 5C). Some strains of *E. coli* are known to induce Th17 cells *in vivo* (Britton et al., 2020; Viladomiu et al., 2017), and IL-23 is a key cytokine that regulates Th17 cell development (Hue et al., 2006). We wondered if genomic sequence similarity between strains of *E. coli* could explain differences in the capacity to elicit IL-23. We found no correlation between whole genome similarity between pairs of *E. coli* strains and their differential ability to induce IL-23 (R^2^ = 0.012, p = 0.23, Pearson correlation; Figure S5E). However, when we looked at the genomes within sub-species of *E. coli* with >65% similarity (Figure S5E), we found that the pairwise genomic similarity over the whole genome negatively correlated with the difference in IL-23 induction (right R^2^ = 0.25, p = 0.007, Pearson correlation; Figure 5F), suggesting that evolution may explain variation in IL-23 induction among closely related strains but does not explain variation between strains that share less sequence homology.

We hypothesized that horizontal gene transfer may disrupt the linkage between whole-genome variation and causal genes in more distantly related genomes. We therefore increased the resolution of our genomic approach to examine the relationship between defined protein-coding regions of the *E. coli* genome and the capacity to elicit IL-23 from myeloid cells. Within a window of 5 consecutive genes moving along the length of the genome (using *E. coli* MG1655 as a reference genome), we calculated the pairwise distance at the protein level and correlated these distances with the pairwise distance in IL-23 induction between strains of *E. coli* (Extended data). Significant associations coalesced around consecutive 5-gene windows at specific genomic coordinates (Figure 5G), consistent with genetic linkage events and the genetic architecture of bacteria where functionally similar genes are often located in close proximity. Among the top 100 strongest correlations (p < 0.0005, Spearman correlation with BH correction), we found enrichments in three different functional gene clusters describing ABC transporters (Keseler et al., 2011), one of which (FunGC 56) is involved in the membrane transport of bacterial peptidoglycans (Figures 5G, S5F). Upon examination of these loci we found significant positive correlations between the pairwise protein distance across consecutive genes at these genomic locations and the pairwise difference in the capacity to induce IL-23 (FunGC 43 ρ = 0.37, p = 2.9×10^-8^; FunGC 52 ρ = 0.28, p < 1×10^-10^; FunGC 56 ρ = 0.43, p < 1×10^-10^; Spearman correlation; Figure 5H, Supplementary Table 4). These data demonstrate a combination of functional and genomic approaches can identify genetic elements that associate with variation in innate immunogenicity between strains of *E. coli.*

## DISCUSSION

Over the past decade there has been great progress towards defining the structure and the composition of the human gut microbiome. Translating these findings into strategies aimed at manipulating the microbiota for therapeutic gain remains a challenge however. The vast number of distinct strains that comprise the human gut microbiota (Faith et al., 2020) implies near-infinite interpersonal variation with unknown physiological consequences to the host. Here, we use a high-throughput screening platform to explore fundamental questions regarding the impact of gut-derived bacterial strains on innate immune phenotypes. We demonstrate that innate immune responses to human gut microbes are reproducible, highly diverse and can be as strong as responses to enteric pathogens. We find that genetics are a strong predictor of the type and magnitude of the innate immune response, from broad enrichments at the phylum level to variation in immunogenicity between individual strains of *E. coli* and *B. ovatus.* Employing novel methods in comparative bacterial genomics, we associate variation in innate immune responses with specific genomic loci.

How recognition of commensal organisms by the innate immune system impacts host physiology is not well characterized. TLR recognition via MyD88 signaling is required for microbiota-dependent immune phenotypes *in vivo,* but how individual components of the microbiota contribute to these responses is not known (Mishima et al., 2019; Mortha et al., 2014; Rakoff-Nahoum et al., 2004). As expected, we find that innate immune cytokine secretion by dendritic cells in response to commensal bacteria depends on MyD88 signaling. Strikingly, myeloid cells differentially rely upon TLR2 and TLR4 to sense microbes from Bacteroidetes and Proteobacteria respectively. Bacterial species that are recognized by TLR2 induce more colonic FoxP3^+^ Treg cells than bacterial species recognized by TLR4. Others have shown that microbial TLR2 ligands are sufficient to induce mucosal FoxP3^+^ Treg cells (Round and Mazmanian, 2010), and that Treg homeostasis is disrupted in mice with MyD88-deficient macrophages (Mortha et al., 2014). Together this suggests the recognition of specific microbes by TLR2 may be a critical regulator of mucosal tolerance. Furthermore, a shift in the ratio of Bacteroidetes and Proteobacteria is a common feature of disease-associated human microbiotas (Contijoch et al., 2019; Gevers et al., 2014), and our data suggests a pathway by which microbiotas with this perturbed ratio may contribute to immune dysregulation in the host.

In addition to variation in innate immune phenotypes across phyla, we also find extraordinary levels of variation in the magnitude of innate responses to species within the same genus, and even more surprisingly, to strains within the same species. TNFα varies across a collection of 10 distinct strains of *B. ovatus,* an *in vitro* phenotype which correlates with the capacity of each strain to induce fecal IgA *in vivo.* The mechanism by which *B. ovatus* induces IgA remains unknown (Yang et al., 2020), and the varied induction of TNFα is likely representative of the complex interactions between distinct bacterial strains and the host.

How the immunologic properties of bacterial strains change following transfer into recipient individuals is not clear. Others have described examples of commensal bacterial phenotypes that change depending on the physiology of the host (Becattini et al., 2021). FMT is widely used to treat recurrent *Clostridioides difficile* infection (Kelly et al., 2016; van Nood et al., 2013), and the stability of immune phenotypes over time is necessary if transmission of bacterial strains as an immunotherapy is to progress (Faith et al., 2015). We find that the cytokine induction to commensal strains remains unchanged when the identical strain is re-isolated from human FMT recipients 11 months following transplant, demonstrating that the immunogenic properties of bacterial strains are stable over time and between hosts.

Particular strains of *E. coli* are known to drive mucosal Th17-biased immune responses (Britton et al., 2020; Viladomiu et al., 2017), but the mechanisms by which these strains elicit Th17 immunity are not known. We find that IL-23 induction varies across 14 distinct strains of *E. coli.* Whole genome similarity correlates with IL-23 induction only between closely related strains of *E. coli.* Using a novel bioinformatics approach, we correlate differences across protein-coding regions with IL-23 induction and identify several associated loci, demonstrating the potential to combine functional and genomic analyses to uncover strain-level mechanisms between the host and the microbiota.

Cross-referencing high-dimensional functional microbiome studies can provide insights into the signals that drive mucosal immune phenotypes. We integrate our study with a published dataset describing mucosal immune cell populations following monocolonization with bacterial species from the human gut microbiota (Geva-Zatorsky et al., 2017). We find that *in vitro* TLR2 stimulation across species correlates with colonic B cell frequencies, consistent with observations that TLR2 signaling helps maintain colonic B cells (Mishima et al., 2019). Notably, we also find that *in vivo* ILC3 development strongly correlates with *in vitro* TLR2 stimulation. Murine ILC3s do not express TLRs, yet these cells are known to translate cues from commensal bacteria via crosstalk with macrophages and dendritic cells (Mortha et al., 2014). Future work will need to investigate the links between TLR2, macrophages and ILCs in the context of specific signals from the microbiota, especially considering the importance of TLR2 stimulation in the development of human ILC3s (Crellin et al., 2010).

An understanding of how PAMPs synergize to boost innate immune responses has implications for vaccine adjuvant design. There are currently very few FDA approved adjuvants, and many consist of only a single individual TLR ligand (Tom et al., 2019). Our data demonstrates that single TLR ligands are not as effective at stimulating immune responses as whole bacteria. Combinations of common PAMPs elicit diverse and tunable innate immune responses, and specific pairs of TLR ligands synergize to boost responses. These combinatorial approaches can be leveraged to uncover immune synergies to create more potent adjuvant formulations.

Like all high-throughput *in vitro* screens, our approach is not without limitations. It is well established that tissue-resident macrophages and dendritic cells have heterogenous phenotypes that differ from bone marrow derived myeloid cells. Future work will need to demonstrate how responses to gut-derived bacteria compare across different sub-populations of macrophages and dendritic cells. Given the large number of strains in our bacterial library and the challenge of defining CFU among diverse bacteria, this *in vitro* screening platform does not use a consistent MOI. However, we find that differences in bacterial concentration (as measured by OD600nm) do not explain the variation of cytokine secreted in response to our bacterial collection. Finally, most of the bacteria in this study are anaerobic and not metabolically active during coculture with macrophages and dendritic cells. Considering our model system measures responses to PAMPs, we believe our measurements are likely unaffected by metabolic inactivity.

In summary, we characterize the immense range of innate immune phenotypes elicited by bacteria derived from the human gut microbiota. We model these responses, identify TLR requirements for the recognition of distinct taxa, and show the immunogenicity of strains is stably maintained across human transmission and across time. Finally, we find numerous cases where *in vitro* immunogenicity is predictive of *in vivo* alterations of immune function, and demonstrate a proof-of-concept comparative genomics approach to identify genetic regions associated with strain-level variation in immunogenicity.

## METHODS

### Mice and gnotobiotic experiments

All animal studies were carried out in accordance with protocols approved by the Institutional Animal Care and Use Committee in Icahn School of Medicine at Mount Sinai. Six to eight-week-old WT C57BL/6J (Jackson Laboratory stock #000664), MyD88^KO^ (Jackson Laboratory stock #009088), TLR2^KO^ (Jackson Laboratory stock #004650), TLR4^KO^ (Jackson Laboratory stock #029015) mice were purchased from Jackson Labs. Mice were kept under specific pathogen-free conditions at the Icahn School of Medicine at Mount Sinai Gnotobiotic Facility. Germ free C57BL/6J were bred in isolators at the Icahn School of Medicine at Mount Sinai Gnotobiotic Facility. Mice were colonized by oral gavage with pure cultures of isolated strains at 6-8 weeks old. Following colonization, gnotobiotic mice were housed in barrier cages under positive pressure and handled using strict aseptic technique. Mice were colonized for 3 weeks before analysis of lamina propria cell populations or fecal IgA concentrations. Some gnotobiotic monocolonization experiments were performed as a part of other published studies to understand relationships with mucosal immune phenotypes (Geva-Zatorsky et al., 2017; Yang et al., 2020).

### Human commensal library construction

Bacterial cultures from human stool samples were previously isolated from de-identified individuals and stored in 96well plates (1 plate per individual; Aggarwala et al., 2021; Britton et al., 2019). To generate our screening library, we regrew culture collections from several different individuals in liquid LYBHIv4 media (37 g/L Brain Heart Infusion [Becton Dickinson], 5 g/L yeast extract [Becton Dickinson], 1 g/L each of D-xylose, D-fructose, D-galactose, cellubiose, maltose, sucrose, 0.5 g/L N-acetylglucosamine, 0.5 g/L L-arabinose, 0.5 g/L L-cysteine, 1 g/L malic acid, 2 g/L sodium sulfate, 0.05% Tween 80, 20 μg/mL menadione, 5 mg/L hemin [as histidine-hemitin], 0.1 M MOPS, pH 7.2) for 48 hours under anaerobic conditions. 320 unique isolates from 17 healthy and IBD donors were selected and re-arrayed into a master library. From this master library, 277 strains faithfully regrew (as determined by a background-subtracted OD600 cutoff of 0.1 and by MALDI-TOF identification). Each time the 320-strain culture collection library or the smaller 54-strain library was regrown, we measured and confirmed growth by OD600nm and identity by MALDI-TOF.

### Preparation of primary cell cultures

Bone marrow derived dendritic cells and macrophages were prepared as previously described (Fordham et al., 2012; Inba et al., 1992). Briefly, femurs were flushed with PBS. Cells were resuspended in complete RPMI (RPMI-1640 (Gibco), 10% heat-inactivated FBS (Gibco), 2mM L-glutamine (Gibco), 25mM HEPES (Gibco), 55nM 2-mercaptoethanol (Gibco), 1mM sodium pyruvate (Gibco), 100U/mL penicillin/streptomycin (Gibco)) and plated in a T75 flask at 37°C for 4 hours to removed adherent cells. To differentiate BMDC, non-adherent bone marrow suspensions were washed and plated at a density of 4×10^5^ cells/mL on 6 well tissue culture treated plates (Corning) in complete RPMI supplemented with 20ng/mL mouse GM-CSF (Peprotech) for 8 days. Differentiated BMDC were harvested, washed with complete RPMI, plated in 96-well plates at a density of 50,000 cells/well and rested overnight before use. To differentiate BMDM, non-adherent bone marrow suspensions were washed, resuspended in growth media (DMEM (Gibco), 10% heat-inactivated FBS, 5% heat-inactivated horse serum (Sigma), 2mM L-glutamine, 55nM 2-mercaptoethanol, 1mM sodium pyruvate, 100U/mL penicillin/streptomycin, 10ng/mL mouse M-CSF (Peprotech), seeded at 2×10^5^ cells/mL and placed in polytetrafluoroethylene bags (Welch Fluorocarbon) at 37°C for 7 days. Differentiated BMDM were harvested, washed with complete RPMI, plated in 96 well plates at a density of 50,000 cells/well and rested for 48 hours before use.

### Commensal screening

Bacterial libraries were centrifuged at 3000g for 10 minutes. Supernatants were removed and passed through a 0.2μm filter (Pall 8019). Bacterial cultures were washed twice with PBS and resuspended in an equal volume of RPMI-1640 supplemented with 0.5mg/mL reduced cysteine (Sigma). BMDM and BMDC were inoculated with 5μL of bacterial cultures or filtered bacterial supernatant. 50μg/mL of gentamicin (Thermo) was added after 1 hour. Supernatants were harvested after 24 hours and stored at −80°C. For the initial screen involving the 277-strain library, we performed three independent experiments and included multiple technical replicate conditions per experiment. For experiments using a 54-member sub-sampled bacterial library, we performed four separate experiments with two technical replicate conditions per experiment such that each isolate was sampled 8 times.

### High-throughput ELISA

Cytokine concentrations were measured in culture supernatants by sandwich ELISA (R&D Systems) according to the manufacturer’s instructions, but with volumes scaled for 384-well format. We assayed TNFα, IL-6, IL-10, IL-23, IL-1β, IL-12, IL-4 and IFNγ. Attempts to measure IL-12, IL-4 and IFNγ were either unsuccessful or not reproducible. There was no significant correlation between responses and the gentamicin mean inhibitory concentration of *E. coli* isolates in our library, excluding the possibility that cytokine induction by Proteobacteria is related to antibiotic-resistance. To accurately capture the dynamic range of IL-6, we used an average across two sample dilutions. Samples were loaded into 384 well MaxiSorp flat-bottom plates (Nunc) using a Biomek FX^P^ liquid handler with a 96-span multichannel pipettor. All ELISA washes were performed using a microplate washer (BioTek 405TS).

### TLR reporter cell lines

HEK-Blue mTLR2 and HEK-Blue mTLR4 cell lines (Invivogen) were maintained following the manufacturer’s recommendations. All assays were performed using endotoxin-free reagents and plasticware. Conditioned media from bacterial cultures were filtered (0.2μm) and diluted in endotoxin-free PBS to create a 5-fold dilution series per sample of 4%, 0.8%, 0.16% and 0.032% (v:v). Half-area tissue-culture-treated 96 well plates were filled with 10μl of each diluted sample followed by 90μl of the reporter cell lines suspended in HEK-Blue Detection Media (Invivogen). Assays were performed with cell densities of 1.4 x 10^5^ cells/well (mTLR2) and 0.7 x 10^5^ cells/well (mTLR4). Assay plates were incubated at 37°C with 5% CO_2_ and the absorbance (600nm) of the colorimetric substrate was measured 16 hours later. The signal from a non-TLR expressing control cell line (HEK-Blue Null-2) was subtracted from the signal of the TLR2 and TLR4 reporter lines. Data presented as either area under the curve (AUC) or the OD600nm at the sample dilution which provided the largest dynamic range.

### Lamina propria cell isolation and flow cytometry

Cells from mouse lamina propria were isolated as previously described (Britton et al., 2019). Briefly, tissues were washed and de-epithelized in HBSS supplemented with 5 mM EDTA, 15 mM HEPES and 5% FBS. Remaining tissue was digested with 0.5 mg/ml Collagenase Type IV (Sigma Aldrich C5138) and 0.5 mg/ml DNase1 (Sigma Aldrich DN25) in HBSS with 5% FBS. Mononuclear cells were obtained by passing through 100 μm and 40 μm strainers. Cells were stained with the following antibodies: CD45 Brilliant Violet (BV)750 (BioLegend), CD3 BV421 (BioLegend), CD4 BV786 (ThermoFisher Scientific), RORgt APC (ThermoFisher Scientific), FoxP3 AlexaFluor488 (ThermoFisher Scientific). Intracellular staining was performed sequentially after surface staining using FoxP3 Fixation/Permeabilization buffers (ThermoFisher Scientific). Dead cells were excluded using a fixable viability dye (eFluor506, ThermoFisher Scientific). Data were acquired on a five laser Aurora Spectral Cytometer (Cytek). Spectral unmixing was performed in Spectraflow (Cytek) and further analysis performed in FlowJo X (BD Bioscience).

### Quantifying pairwise kmer distances

The kmer overlap between each pair of E. coli genomes, A and B, were determined by generating a hash for genome A with kmer size 20 and quantifying the proportion of kmers shared in both genomes A and B divided by the total number of kmers in A. These distances were independently calculated in both directions, and the mean distance was used.

### Comparative bacterial genomics

*E. coli* genomes were annotated with prokka version 1.12 (Seemann, 2014) and orthologs were determined with OrthoFinder version 2.5.4 (Emms and Kelly, 2019). The *E. coli* strain K-12 substrain MG1655 (Blattner et al., 1997) was used as a reference to determine local gene order. We calculated a 5-gene moving average of the protein distance along the length of the genome using a Clustal Omega alignment for all orthologs in a given window (i.e. for a set of 5 consecutive genes we used the protein distance for all genes with an ortholog across the *E. coli* strains in our collection) (Sievers et al., 2011). Functional clusters were obtained from EcoCyc (Keseler et al., 2011). Functional clusters which contained genes with no corresponding ortholog across our E. coli collection were excluded. Finally, for visualization purposes, we calculated all possible moving gene-windows of size 1 to 10 for the three highlighted genetic regions that overlapped with enriched functional clusters (Figure 5H) in order to identify the best-fit region containing correlated genes.

### Data integration, statistical analysis, modelling and plotting

Data on immune cell population frequencies in monocolonized mice were obtained from previously published work (Geva-Zatorsky et al., 2017). The dataset included 21 species that were also present in the screening library used in this study. In cases where either dataset contained multiple strains of the same species, the mean data from all strains was used for comparative analyses. Analyses throughout this study were performed using RStudio v1.1.463. All p values less than 1×10^-10^ were rounded. Heatmaps were constructed using either *ComplexHeatmap* (Gu et al., 2016) or ggplot2. For plots depicting fold-change of knockout responses relative to wildtype, the bottom 3% of responses per analyte were thresholded for visualization purposes. K-nearest neighbor classification was performed using the R package *class* (k=15). Stepwise regression (using AIC minimization) was applied using R package *MASS.* Procedures for fitting elastic net regression (λ selected using 10-fold cross validation) were performed with the R package *glmnet.* The multiple linear regression for modelling responses to TLR ligand combinations including interaction terms is:

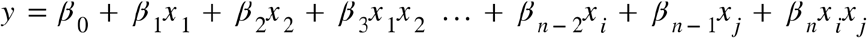

Where x_1_ = TLR ligand 1, x_2_ = TLR ligand 2 …, x_i_ = TLR ligand *i, x_j_* = TLR ligand *j, n* = number of terms.

## Acknowledgements

This work was supported in part by the staff and resources of the Mount Sinai Gnotobiotic Facility, the Mount Sinai Flow Cytometry Core, and the Scientific Computing Division at the Icahn School of Medicine at Mount Sinai. Whole-genome assembled sequences (FASTA) of strains have been deposited under project number PRJNA637878. We thank C. Fermin, E. Vazquez, and G.N. Escano for gnotobiotic husbandry and for technical support, V. Aggarwala for help with bacterial genomics. This work was supported by National Institutes of Health grants (nos. NIDDK DK112978, NIDDK DK124133, NIDDK DK123749), an NIH F30 to M.P.S. (DK124978), and a SUCCESS philanthropic award and a Crohn’s and Colitis Foundation RFA award to G.J.B. (no. 580924) and J.J.F. (nos. 632758, 651867).

## Author contributions

M.P.S., G.J.B and J.J.F. wrote the manuscript. M.P.S., S.S.S., G.J.B., Z.L., I.M. collected the samples and performed the experiments. M.P.S., S.S.S., C.Y., S.M., G.J.B., J.J.F. analyzed and interpreted the data. All authors read, provided critical feedback and approved the final manuscript.

---

**Figure S1.**
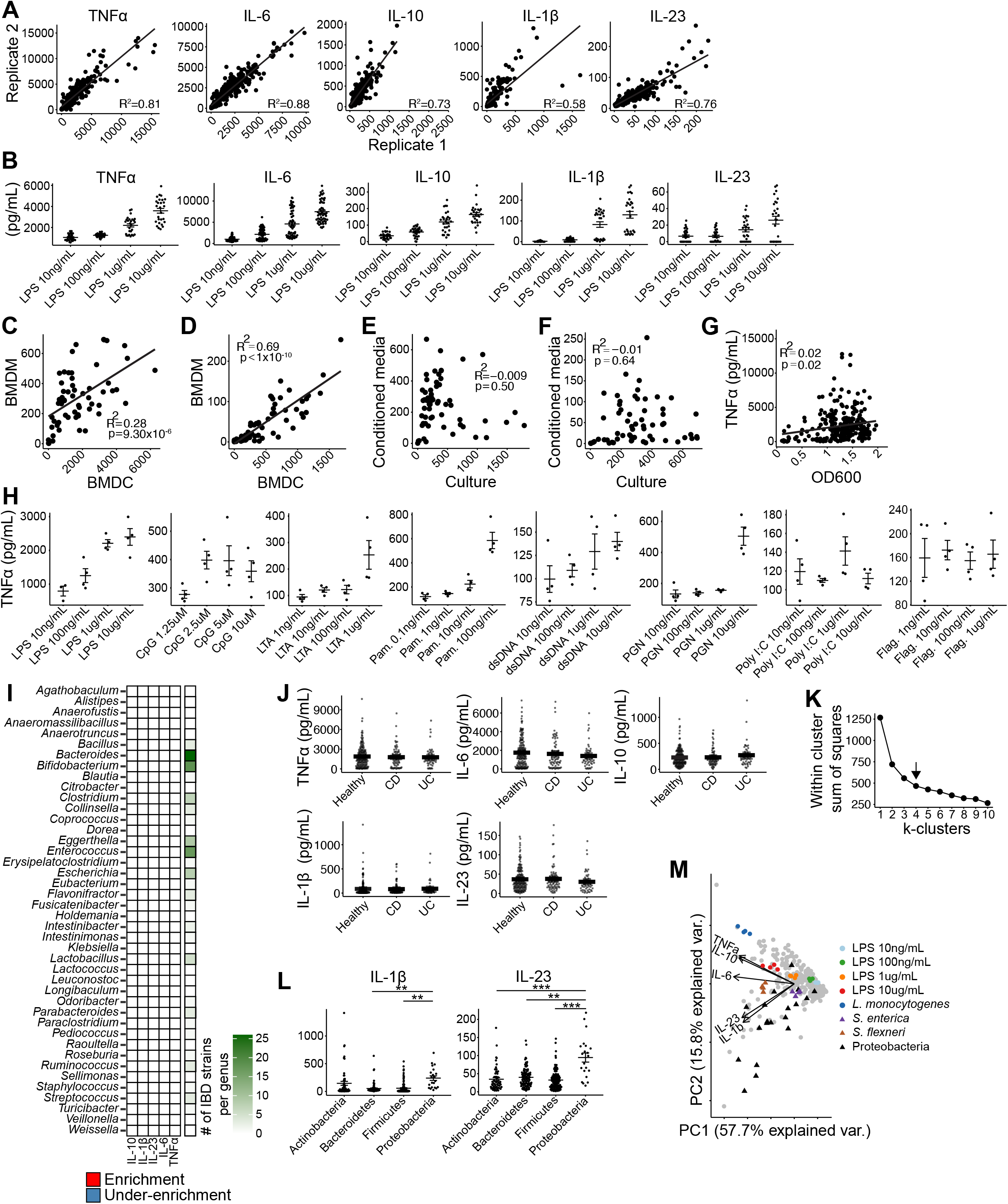
(A) Biologic replicates performed at different times correlate. (B) LPS dose-response relationships (C) TNFα secretion by BMDC versus BMDM stimulated with live bacterial cultures correlates. (D) TNFα secretion by BMDC versus BMDM stimulated with bacterial conditioned media correlates. (E) TNFα secretion by BMDC stimulated with bacterial conditioned media versus bacterial cultures does not correlate. (F) TNFα secretion by BMDM stimulated with bacterial conditioned media versus bacterial cultures does not correlate. (G) TNFα secretion by BMDC weakly correlates with OD600 measurements taken during stationary phase. (H) Dose-dependent relationships to the indicated TLR ligands. (I) Enrichment for cytokine stimulation based on IBD status per genus. Enrichment with a two-sided Fisher’s exact test using IBD-association and membership above the median response as the contingencies. (J) Cytokine induction by isolates derived from healthy donors, Crohn’s disease patients (CD) or Ulcerative Colitis patients (UC). Solid horizontal lines indicate mean ± SEM. Each point represents the mean response of an individual isolate. (K) Sum of squares distance to nearest cluster center for the data in Figure 1D for values of k 1 through 10. Arrow represents chosen value for k. (L) IL-1β and IL-23 secretion by BDMCs in response to strains from the indicated phylum. Solid horizontal lines indicate mean ± SEM. * p < 0.05, ** p < 0.01, *** p < 0.001. Students t-test with BH correction. Each point represents the mean response of an individual isolate. (M) Principal component analysis of innate cytokine profiles. Linear regression p values in (A, C-G) calculated by F-test. dsDNA, double-stranded genomic DNA from E. coli K12. Flag, flagellin. LTA, lipoteichoic acid. Pam, Pam3CSK4. PGN, peptidoglycan.

**Figure S2.**
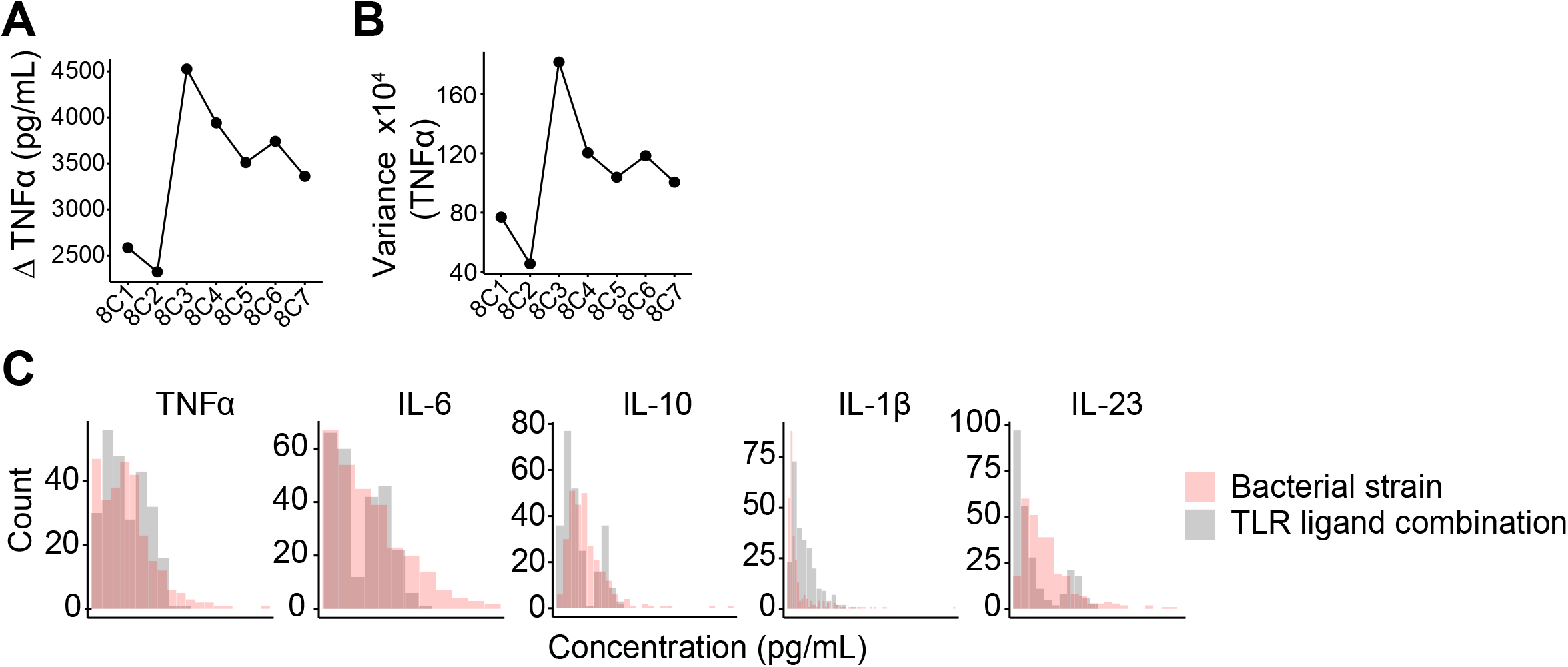
(A) Range of TNFα response as a function of *K*. Minimum value was subtracted from maximum value for each set of combinations. (B) Variance of TNFα response as a function of *K*. (C) Histogram of responses to 277 bacterial strains and to 255 TLR ligand combinations. Bin width: *h*=2×IQR×*n*^-1/3^

**Figure S3.**
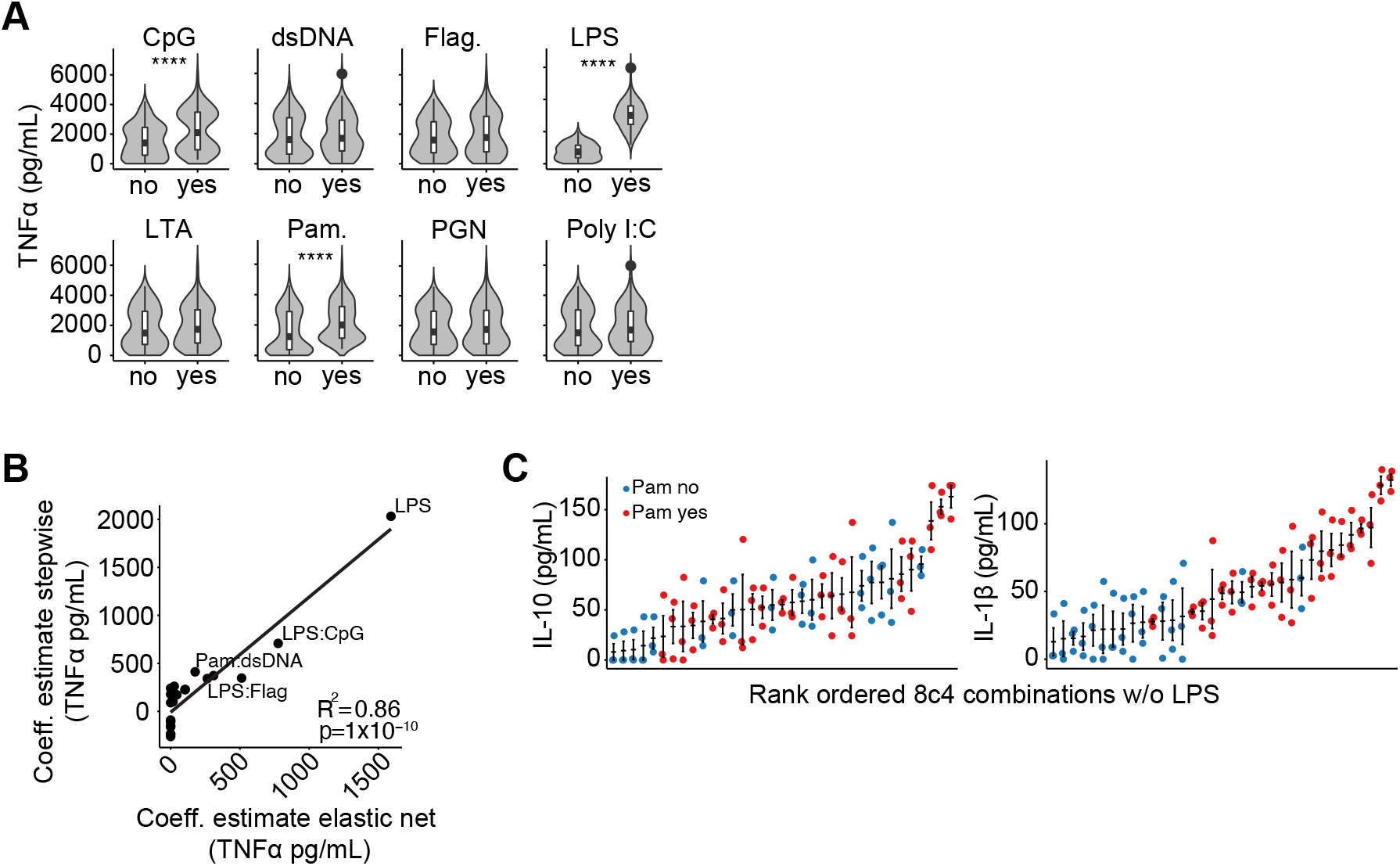
(A) TNFα responses in the presence or absence of the indicated TLR ligands. **** p < 0.0001. Student’s t-test. (B) Model parameter estimates determined using stepwise regression correlates with estimates determined using elastic net regression. Certain significant pairwise synergistic interactions are labeled. Regression p-values calculated by F-test. (C) Rank ordered IL-10 and IL-1β responses to TLR ligand combinations of 8 choose 4 excluding conditions containing LPS.

**Figure S4.**
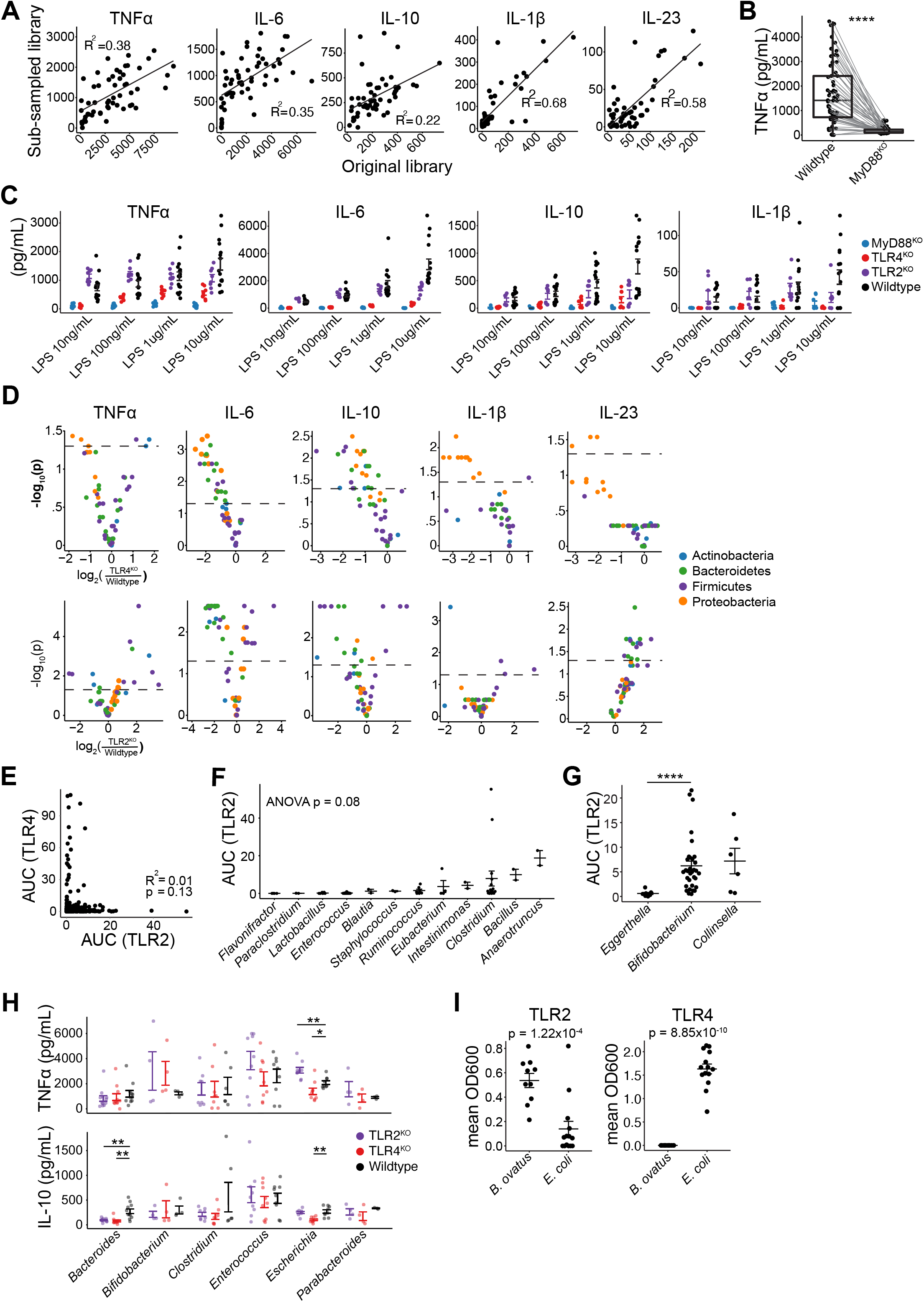
(A) BMDC responses using the original 277-member commensal library and the subsampled library. Regression p-value calculated by F-test. (B) Wildtype versus MyD88^KO^ BMDC responses to 54 different gut-derived bacterial strains. p < 0.0001. Paired student’s t-test. (C) BMDC dose-response to LPS. (D) Volcano plots showing fold change from wildtype of TLR2^KO^ and TLR4^KO^ BMDC. A threshold was applied to very low measurements as described in the methods. (E) TLR2 and TLR4 stimulation by gut derived bacteria are non-overlapping. (F) The majority of genera within Firmicutes do not stimulate TLR2. (G) Specific genera within Actinobacteria more potently stimulate TLR2 than others. (H) TNFα and IL-10 responses across indicated knockout BMDC cell lines. Each point represents the mean response to a particular isolate. * p< 0.05, ** p < 0.01. Student’s t-test with BH correction. (I) *B. ovatus* and *E. coli* differentially rely on TLR2 and TLR4. Conditioned media from 10 distinct strains of *B. ovatus* and 14 distinct strains of *E. coli* were used. Student’s t-test.

**Figure S5.**
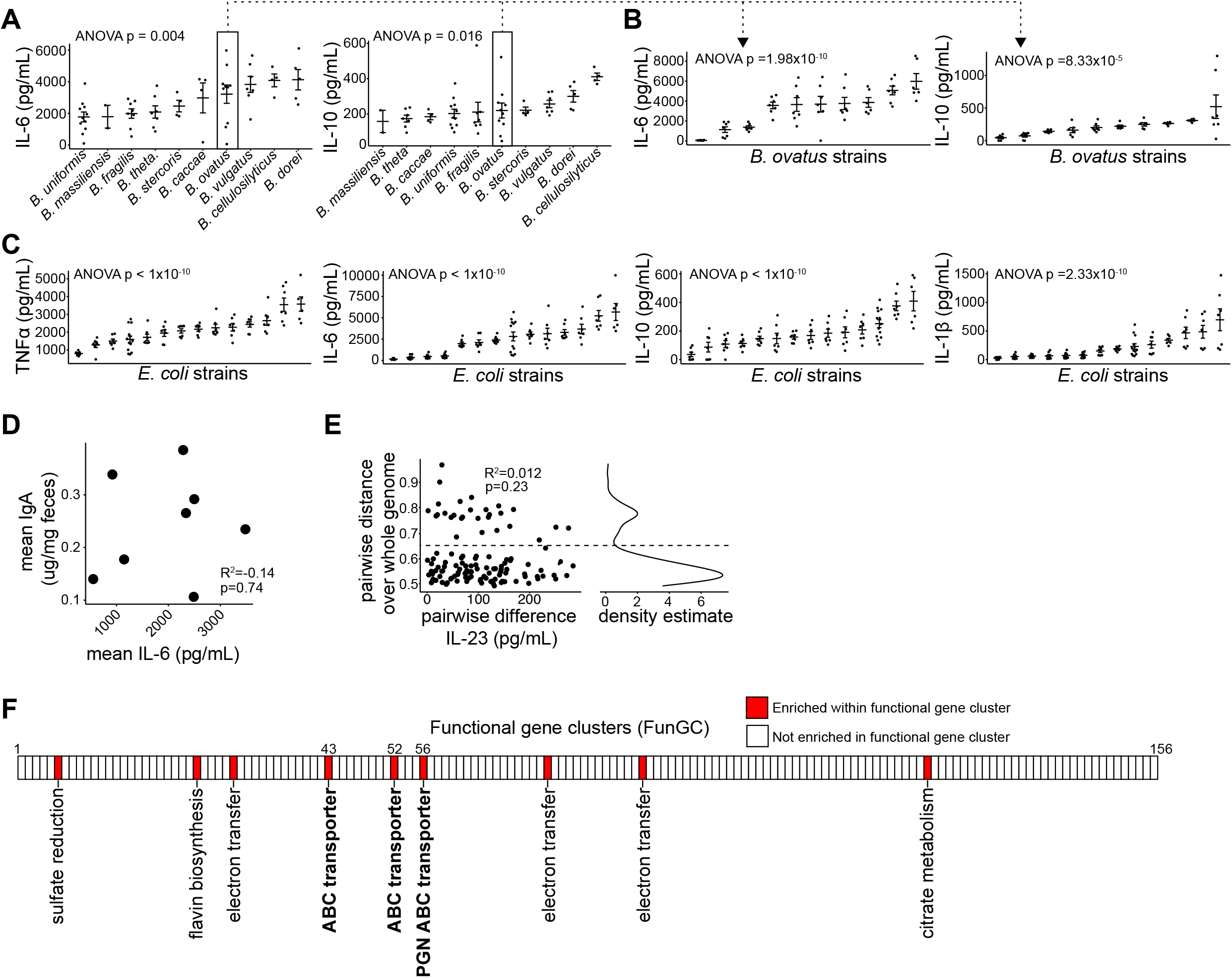
(A-B) IL-6 and IL-10 induction varies across species of *Bacteroides* and across strains of *B. ovatus.* Species level plot each point represents the mean response of an individual isolate. Strain level plot each point represents a replicate condition. (C) TNFα, IL-6, IL-10 and IL-1β induction varies across strains of *E. coli.* Each point represents a replicate condition. (D) IL-6 secretion by BMDC in response to *B. ovatus* strains versus fecal IgA measurements from gnotobiotic mice monocolonized with corresponding *B. ovatus* strains. Mean IL-6 measurements represent the average of four biologic replicates. Mean IgA measurements represent an average from 4-8 mice per strain. Regression p values calculated by F-test. (E) Left: Pairwise distance across the whole genome versus the pairwise difference in IL-23 induction between *E. coli* strains. Regression p values calculated by F-test. Right: Density estimation of pairwise distances across *E. coli* genomes. Dotted line represents 65% sequence homology and delineates an *E. coli* sub-species. (F) Enrichment for *E. coli* genes associating strongly with IL-23 induction within 156 defined functional gene clusters (EcoCyc; Keseler et al., 2011). Enrichment calculated using a two-sided Fisher’s exact test with the presence of a gene associating strongly with IL-23 induction (defined as p < 0.0005, Spearman correlation with BH correction) and the presence within a functional gene cluster as the two contingencies. The 9 FunGCs significantly enriched for genes associating strongly with IL-23 induction are labeled.

## REFERENCES

Abrams, J., Figdor, C.G., De Waal Malefyt, R., Bennett, B., De Vries, J.E., 1991. Interleukin 10(IL-10) inhibits cytokine synthesis by human monocytes: An autoregulatory role of IL-10 produced by monocytes. J. Exp. Med. 174, 1209–1220. https://doi.org/10.1084/jem.174.5.1209

Aggarwala, V., Mogno, I., Li, Z., Yang, C., Britton, G.J., Chen-Liaw, A., Mitcham, J., Bongers, G., Gevers, D., Clemente, J.C., Colombel, J.F., Grinspan, A., Faith, J., 2021. Precise quantification of bacterial strains after fecal microbiota transplantation delineates long-term engraftment and explains outcomes. Nat. Microbiol. 6, 1309–1318. https://doi.org/10.1038/s41564-021-00966-0

Aleyas, A.G., George, J.A., Han, Y.W., Rahman, M.M., Kim, S.J., Han, S.B., Kim, B.S., Kim, K., Eo, S.K., 2009. Functional Modulation of Dendritic Cells and Macrophages by Japanese Encephalitis Virus through MyD88 Adaptor Molecule-Dependent and - Independent Pathways. J. Immunol. 183, 2462–2474. https://doi.org/10.4049/JIMMUNOL.0801952

Atarashi, K., Tanoue, T., Ando, M., Kamada, N., Nagano, Y., Narushima, S., Suda, W., Imaoka, A., Setoyama, H., Nagamori, T., Ishikawa, E., Shima, T., Hara, T., Kado, S., Jinnohara, T., Ohno, H., Kondo, T., Toyooka, K., Watanabe, E., Yokoyama, S.I., Tokoro, S., Mori, H., Noguchi, Y., Morita, H., Ivanov, I.I., Sugiyama, T., Nuñez, G., Camp, J.G., Hattori, M., Umesaki, Y., Honda, K., 2015. Th17 Cell Induction by Adhesion of Microbes to Intestinal Epithelial Cells. Cell. https://doi.org/10.1016/j.cell.2015.08.058

Bafica, A., Santiago, H.C., Goldszmid, R., Ropert, C., Gazzinelli, R.T., Sher, A., 2006. TLR9 and TLR2 Signaling Together Account for MyD88-Dependent Control of Parasitemia in Trypanosoma cruzi Infection. J. Immunol. 177, 3515–3519. https://doi.org/10.4049/jimmunol.177.6.3515

Bafica, A., Scanga, C.A., Feng, C.G., Leifer, C., Cheever, A., Sher, A., 2005. TLR9 regulates Th1 responses and cooperates with TLR2 in mediating optimal resistance to Mycobacterium tuberculosis. J. Exp. Med. Artic. 1715 JEM 202, 1715–1724. https://doi.org/10.1084/jem.20051782

Becattini, S., Sorbara, M.T., Kim, S.G., Littmann, E.L., Dong, Q., Walsh, G., Wright, R., Amoretti, L., Fontana, E., Hohl, T.M., Pamer, E.G., 2021. Rapid transcriptional and metabolic adaptation of intestinal microbes to host immune activation. Cell Host Microbe 29, 378–393.e5. https://doi.org/10.1016/j.chom.2021.01.003

Britton, G.J., Contijoch, E.J., Mogno, I., Vennaro, O.H., Llewellyn, S.R., Ng, R., Li, Z., Mortha, A., Merad, M., Das, A., Gevers, D., McGovern, D.P.B., Singh, N., Braun, J., Jacobs, J.P., Clemente, J.C., Grinspan, A., Sands, B.E., Colombel, J.F., Dubinsky, M.C., Faith, J.J., 2019. Microbiotas from Humans with Inflammatory Bowel Disease Alter the Balance of Gut Th17 and RORγt + Regulatory T Cells and Exacerbate Colitis in Mice. Immunity. https://doi.org/10.1016/j.immuni.2018.12.015

Britton, G.J., Contijoch, E.J., Spindler, M.P., Aggarwala, V., Dogan, B., Bongers, G., Mateo, L.S., Baltus, A., Das, A., Gevers, D., Borody, T.J., Kaakoush, N.O., Kamm, M.A., Mitchell, H., Paramsothy, S., Clemente, J.C., Colombel, J.F., Simpson, K.W., Dubinsky, M.C., Grinspan, A., Faith, J.J., 2020. Defined microbiota transplant restores Th17/RORγt+ regulatory T cell balance in mice colonized with inflammatory bowel disease microbiotas. Proc. Natl. Acad. Sci. U. S. A. 117, 21536–21545. https://doi.org/10.1073/pnas.1922189117

Britton, G.J., Faith, J.J., 2021. Causative Microbes in Host-Microbiome Interactions. Annu. Rev. Microbiol. 75, 223–242. https://doi.org/10.1146/annurev-micro-041321-042402

Contijoch, E.J., Britton, G.J., Yang, C., Mogno, I., Li, Z., Ng, R., Llewellyn, S.R., Hira, S., Johnson, C., Rabinowitz, K.M., Barkan, R., Dotan, I., Hirten, R.P., Fu, S.C., Luo, Y., Yang, N., Luong, T., Labrias, P.R., Lira, S., Peter, I., Grinspan, A., Clemente, J.C., Kosoy, R., Kim-Schulze, S., Qin, X., Castillo, A., Hurley, A., Atreja, A., Rogers, J., Fasihuddin, F., Saliaj, M., Nolan, A., Reyes-Mercedes, P., Rodriguez, C., Aly, S., Santa-Cruz, K., Peters, L., Suárez-Fariñas, M., Huang, R., Hao, K., Zhu, J., Zhang, B., Losic, B., Irizar, H., Song, W.M., Di Narzo, A., Wang, W., Cohen, B.L., DiMaio, C., Greenwald, D., Itzkowitz, S., Lucas, A., Marion, J., Maser, E., Ungaro, R., Naymagon, S., Novak, J., Shah, B., Ullman, T., Rubin, P., George, J., Legnani, P., Telesco, S.E., Friedman, J.R., Brodmerkel, C., Plevy, S., Cho, J.H., Colombel, J.F., Schadt, E.E., Argmann, C., Dubinsky, M., Kasarskis, A., Sands, B., Faith, J.J., 2019. Gut microbiota density influences host physiology and is shaped by host and microbial factors. Elife 8. https://doi.org/10.7554/eLife.40553

Crellin, N.K., Trifari, S., Kaplan, C.D., Satoh-Takayama, N., Di Santo, J.P., Spits, H., 2010. Regulation of cytokine secretion in human CD127+ LTi-like innate lymphoid cells by toll-like receptor 2. Immunity 33, 752–764. https://doi.org/10.1016/j.immuni.2010.10.012

De Nardo, D., De Nardo, C.M., Nguyen, T., Hamilton, J.A., Scholz, G.M., 2009. Signaling Crosstalk during Sequential TLR4 and TLR9 Activation Amplifies the Inflammatory Response of Mouse Macrophages. J. Immunol. 183, 8110–8118. https://doi.org/10.4049/jimmunol.0901031

Eckburg, P.B., Bik, E.M., Bernstein, C.N., Purdom, E., Dethlefsen, L., Sargent, M., Gill, S.R., Nelson, K.E., Relman, D.A., 2005. Diversity of the human intestinal microbial flora, Science. https://doi.org/10.1126/science.1110591

Faith, J.J., Chen-Liaw, A., Aggarwala, V., Kaakoush, N.O., Borody, J., Mitchell, H., Kamm, M.A., Paramsothy, S., Snitkin, E.S., Mogno, I., 2020. Strain population structure varies widely across bacterial species and predicts strain colonization in unrelated individuals. bioRxiv 2020.10.17.343640. https://doi.org/10.1101/2020.10.17.343640

Faith, J.J., Colombel, J.F., Gordon, J.I., 2015. Identifying strains that contribute to complex diseases through the study of microbial inheritance. Proc. Natl. Acad. Sci. U. S. A. https://doi.org/10.1073/pnas.1418781112

Faith, J.J., Guruge, J.L., Charbonneau, M., Subramanian, S., Seedorf, H., Goodman, A.L., Clemente, J.C., Knight, R., Heath, A.C., Leibel, R.L., Rosenbaum, M., Gordon, J.I., 2013. The long-term stability of the human gut microbiota. Science (80-.). 341. https://doi.org/10.1126/science.1237439

Feagan, B.G., Sandborn, W.J., Gasink, C., Jacobstein, D., Lang, Y., Friedman, J.R., Blank, M.A., Johanns, J., Gao, L.-L., Miao, Y., Adedokun, O.J., Sands, B.E., Hanauer, S.B., Vermeire, S., Targan, S., Ghosh, S., de Villiers, W.J., Colombel, J.-F., Tulassay, Z., Seidler, U., Salzberg, B.A., Desreumaux, P., Lee, S.D., Loftus, E. V., Dieleman, L.A., Katz, S., Rutgeerts, P., 2016. Ustekinumab as Induction and Maintenance Therapy for Crohn’s Disease. N. Engl. J. Med. https://doi.org/10.1056/NEJMoa1602773

Fordham, J.B., Hua, J., Morwood, S.R., Schewitz-Bowers, L.P., Copland, D.A., Dick, A.D., Nicholson, L.B., 2012. Environmental conditioning in the control of macrophage thrombospondin-1 production. Sci. Rep. 2. https://doi.org/10.1038/srep00512

Geva-Zatorsky, N., Sefik, E., Kua, L., Pasman, L., Tan, T.G., Ortiz-Lopez, A., Yanortsang, T.B., Yang, L., Jupp, R., Mathis, D., Benoist, C., Kasper, D.L., 2017. Mining the Human Gut Microbiota for Immunomodulatory Organisms. Cell 168, 928–943.e11. https://doi.org/10.1016/j.cell.2017.01.022

Gevers, D., Kugathasan, S., Denson, L.A., Vázquez-Baeza, Y., Van Treuren, W., Ren, B., Schwager, E., Knights, D., Song, S.J., Yassour, M., Morgan, X.C., Kostic, A.D., Luo, C., González, A., McDonald, D., Haberman, Y., Walters, T., Baker, S., Rosh, J., Stephens, M., Heyman, M., Markowitz, J., Baldassano, R., Griffiths, A., Sylvester, F., Mack, D., Kim, S., Crandall, W., Hyams, J., Huttenhower, C., Knight, R., Xavier, R.J., 2014. The treatment-naive microbiome in new-onset Crohn’s disease. Cell Host Microbe 15, 382–392. https://doi.org/10.1016/j.chom.2014.02.005

Gu, Z., Eils, R., Schlesner, M., 2016. Complex heatmaps reveal patterns and correlations in multidimensional genomic data. Bioinformatics 32, 2847–2849. https://doi.org/10.1093/BIOINFORMATICS/BTW313

Hou, B., Reizis, B., DeFranco, A., 2008. Toll-like receptors activate innate and adaptive immunity by using dendritic cell-intrinsic and -extrinsic mechanisms. Immunity 29, 272–282. https://doi.org/10.1016/J.IMMUNI.2008.05.016

Hue, S., Ahern, P., Buonocore, S., Kullberg, M.C., Cua, D.J., McKenzie, B.S., Powrie, F., Maloy, K.J., 2006. Interleukin-23 drives innate and T cell–mediated intestinal inflammation. J. Exp. Med. https://doi.org/10.1084/jem.20061099

Hussain, S., Johnson, C.G., Sciurba, J., Meng, X., Stober, V.P., Liu, C., Cyphert-Daly, J. M., Bulek, K., Qian, W., Solis, A., Sakamachi, Y., Trempus, C.S., Aloor, J.J., Gowdy, K.M., Foster, W.M., Hollingsworth, J.W., Tighe, R.M., Li, X., Fessler, M.B., Garantziotis, S., 2020. Tlr5 participates in the TLR4 receptor complex and promotes MyD88-dependent signaling in environmental lung injury. Elife 9. https://doi.org/10.7554/ELIFE.50458

Inba, K., Inaba, M., Romani, N., Aya, H., Deguchi, M., Ikehara, S., Muramatsu, S., Steinman, R.M., 1992. Generation of large numbers of dendritic cells from mouse bone marrow cultures supplemented with granulocyte/macrophage colonystimulating factor, Journal of Experimental Medicine. https://doi.org/10.1084/jem.176.6.1693

Ivanov, I., Atarashi, K., Manel, N., Brodie, E.L., Shima, T., Karaoz, U., Wei, D., Goldfarb, K.C., Santee, C.A., Lynch, S. V., Tanoue, T., Akemi, I., Itoh, K., Takeda, K., Umesaki, Y., Honda, K., Littman, D.R., 2009. Induction of Intestinal Th17 Cells by Segmented Filamentous Bacteria. Cell 139, 485–498.

Iwasaki, A., Medzhitov, R., 2010. Regulation of adaptive immunity by the innate immune system. Science (80-.). 327, 291–295. https://doi.org/10.1126/science.1183021

Janeway, C.A., Medzhitov, R., 2002. Innate immune recognition. Annu. Rev. Immunol. https://doi.org/10.1146/annurev.immunol.20.083001.084359

Janeway, C.A.J., 1989. Approaching the asymptote? Evolution and revolution in immunology. Cold Spring Harb. Quant. Biol. 54, 1–13.

Jang, S., Uematsu, S., Akira, S., Salgame, P., 2004. IL-6 and IL-10 Induction from Dendritic Cells in Response to Mycobacterium tuberculosis Is Predominantly Dependent on TLR2-Mediated Recognition. J. Immunol. 173, 3392–3397. https://doi.org/10.4049/JIMMUNOL.173.5.3392

Kelly, C.R., Khoruts, A., Staley, C., Sadowsky, M.J., Abd, M., Alani, M., Bakow, B., Curran, P., McKenney, J., Tisch, A., Reinert, S.E., MacHan, J.T., Brandt, L.J., 2016. Effect of fecal microbiota transplantation on recurrence in multiply recurrent clostridium difficile infection a randomized trial. Ann. Intern. Med. 165, 609–616. https://doi.org/10.7326/M16-0271

Keseler, I.M., Collado-Vides, J., Santos-Zavaleta, A., Peralta-Gil, M., Gama-Castro, S., Muniz-Rascado, L., Bonavides-Martinez, C., Paley, S., Krummenacker, M., Altman, T., Kaipa, P., Spaulding, A., Pacheco, J., Latendresse, M., Fulcher, C., Sarker, M., Shearer, A.G., Mackie, A., Paulsen, I., Gunsalus, R.P., Karp, P.D., 2011. EcoCyc: A comprehensive database of Escherichia coli biology. Nucleic Acids Res. 39. https://doi.org/10.1093/nar/gkq1143

Kühn, R., Löhler, J., Rennick, D., Rajewsky, K., Müller, W., 1993. Interleukin-10-deficient mice develop chronic enterocolitis. Cell 75, 263–274. https://doi.org/10.1016/0092-8674(93)80068-P

Lee, Y.N., Lee, Y.T., Kim, M.C., Hwang, H.S., Lee, J.S., Kim, K.H., Kang, S.M., 2014. Fc receptor is not required for inducing antibodies but plays a critical role in conferring protection after influenza M2 vaccination. Immunology 143, 300–309. https://doi.org/10.1111/IMM.12310

Ligumsky, M., Simon, P.L., Karmeli, F., Rachmilewitz, D., 1990. Role of interleukin 1 in inflammatory bowel disease-enhanced production during active disease. Gut 31, 686–689. https://doi.org/10.1136/gut.31.6.686

Long, E.M., Millen, B., Kubes, P., Robbins, S.M., 2009. Lipoteichoic acid induces unique inflammatory responses when compared to other toll-like receptor 2 ligands. PLoS One 4. https://doi.org/10.1371/journal.pone.0005601

Mazmanian, S.K., Cui, H.L., Tzianabos, A.O., Kasper, D.L., 2005. An immunomodulatory molecule of symbiotic bacteria directs maturation of the host immune system. Cell 122, 107–118. https://doi.org/10.1016/j.cell.2005.05.007

Mishima, Y., Oka, A., Liu, B., Herzog, J.W., Eun, C.S., Fan, T.J., Bulik-Sullivan, E., Carroll, I.M., Hansen, J.J., Chen, L., Wilson, J.E., Fisher, N.C., Ting, J.P.Y., Nochi, T., Wahl, A., Victor Garcia, J., Karp, C.L., Balfour Sartor, R., 2019. Microbiota maintain colonic homeostasis by activating TLR2/MyD88/PI3K signaling in IL-10-producing regulatory B cells. J. Clin. Invest. 129, 3702–3716. https://doi.org/10.1172/JCI93820

Mortha, A., Chudnovskiy, A., Hashimoto, D., Bogunovic, M., Spencer, S.P., Belkaid, Y., Merad, M., 2014. Microbiota-dependent crosstalk between macrophages and ILC3 promotes intestinal homeostasis. Science (80-.). 343. https://doi.org/10.1126/science.1249288

Mudter, J., Neurath, M.F., 2007. IL-6 signaling in inflammatory bowel disease: Pathophysiological role and clinical relevance. Inflamm. Bowel Dis. https://doi.org/10.1002/ibd.20148

Napolitani, G., Rinaldi, A., Bertoni, F., Sallusto, F., Lanzavecchia, A., 2005. Selected Toll-like receptor agonist combinations synergistically trigger a T helper type 1-polarizing program in dendritic cells. Nat. Immunol. 6. https://doi.org/10.1038/ni1223

Netea, M.G., Sutmuller, R., Hermann, C., Van der Graaf, C.A.A., Van der Meer, J.W.M., van Krieken, J.H., Hartung, T., Adema, G., Kullberg, B.J., 2004. Toll-Like Receptor 2 Suppresses Immunity against Candida albicans through Induction of IL-10 and Regulatory T Cells. J. Immunol. 172, 3712–3718. https://doi.org/10.4049/jimmunol.172.6.3712

Olm, M.R., Crits-Christoph, A., Bouma-Gregson, K., Firek, B.A., Morowitz, M.J., Banfield, J.F., 2021. inStrain profiles population microdiversity from metagenomic data and sensitively detects shared microbial strains. Nat. Biotechnol. 39, 727–736. https://doi.org/10.1038/s41587-020-00797-0

Onderdonk, A.B., Bronson, R., Cisneros, R., 1987. Comparison of Bacteroides vulgatus strains in the enhancement of experimental ulcerative colitis. Infect. Immun. 55, 835–836. https://doi.org/10.1128/iai.55.3.835-836.1987

Ozinsky, A., Underhill, D.M., Fontenot, J.D., Hajjar, A.M., Smith, K.D., Wilson, C.B., Schroeder, L., Aderem, A., 2000. The repertoire for pattern recognition of pathogens by the innate immune system is defined by cooperation between Tolllike receptors.

Qin, J., Li, R., Raes, J., Arumugam, M., Burgdorf, K.S., Manichanh, C., Nielsen, T., Pons, N., Levenez, F., Yamada, T., Mende, D.R., Li, J., Xu, J., Li, Shaochuan, Li, D., Cao, J., Wang, B., Liang, H., Zheng, H., Xie, Y., Tap, J., Lepage, P., Bertalan, M., Batto, J.M., Hansen, T., Le Paslier, D., Linneberg, A., Nielsen, H.B., Pelletier, E., Renault, P., Sicheritz-Ponten, T., Turner, K., Zhu, H., Yu, C., Li, Shengting, Jian, M., Zhou, Y., Li, Y., Zhang, X., Li, Songgang, Qin, N., Yang, H., Wang, Jian, Brunak, S., Doré, J., Guarner, F., Kristiansen, K., Pedersen, O., Parkhill, J., Weissenbach, J., Bork, P., Ehrlich, S.D., Wang, Jun, Antolin, M., Artiguenave, F., Blottiere, H., Borruel, N., Bruls, T., Casellas, F., Chervaux, C., Cultrone, A., Delorme, C., Denariaz, G., Dervyn, R., Forte, M., Friss, C., Van De Guchte, M., Guedon, E., Haimet, F., Jamet, A., Juste, C., Kaci, G., Kleerebezem, M., Knol, J., Kristensen, M., Layec, S., Le Roux, K., Leclerc, M., Maguin, E., Melo Minardi, R., Oozeer, R., Rescigno, M., Sanchez, N., Tims, S., Torrejon, T., Varela, E., De Vos, W., Winogradsky, Y., Zoetendal, E., 2010. A human gut microbial gene catalogue established by metagenomic sequencing. Nature 464, 59–65. https://doi.org/10.1038/nature08821

Rakoff-Nahoum, S., Paglino, J., Eslami-Varzaneh, F., Edberg, S., Medzhitov, R., 2004. Recognition of commensal microflora by toll-like receptors is required for intestinal homeostasis. Cell 118, 229–241. https://doi.org/10.1016/J.CELL.2004.07.002

Rath, H.C., Herfarth, H.H., Ikeda, J.S., Grenther, W.B., Hamm, T.E., Balish, E., Taurog, J.D., Hammer, R.E., Wilson, K.H., Sartor, R.B., 1996. Normal Bacteria Stimulate Colitis, Gastritis, and Arthritis in B27 Transgenic Rats Normal Luminal Bacteria, Especially Bacteroides Species, Mediate Chronic Colitis, Gastritis, and Arthritis in HLA-B27/Human 2 Microglobulin Transgenic Rats, The Journal of Clinical Investigation.

Rath, H.C., Wilson, K.H., Sartor, R.B., 1999. Differential induction of colitis and gastritis in HLA-B27 transgenic rats selectively colonized with Bacteroides vulgatus or Escherichia coli. Infect. Immun. 67, 2969–2974. https://doi.org/10.1128/iai.67.6.2969-2974.1999

Reis E Sousa, C., 2004. Toll-like receptors and dendritic cells: for whom the bug tolls. Semin. Immunol. 16, 27–34. https://doi.org/10.1016/J.SMIM.2003.10.004

Round, J.L., Mazmanian, S.K., 2010. Inducible Foxp3+ regulatory T-cell development by a commensal bacterium of the intestinal microbiota. Proc. Natl. Acad. Sci. U. S. A. 107, 12204–12209. https://doi.org/10.1073/pnas.0909122107

Schloss, P.D., Iverson, K.D., Petrosino, J.F., Schloss, S.J., 2014. The dynamics of a family’s gut microbiota reveal variations on a theme, Microbiome. https://doi.org/10.1186/2049-2618-2-25

Seo, S.U., Kamada, N., Muñoz-Planillo, R., Kim, Y.G., Kim, D., Koizumi, Y., Hasegawa, M., Himpsl, S.D., Browne, H.P., Lawley, T.D., Mobley, H.L.T., Inohara, N., Núñez, G., 2015. Distinct Commensals Induce Interleukin-1β via NLRP3 Inflammasome in Inflammatory Monocytes to Promote Intestinal Inflammation in Response to Injury. Immunity 42, 744–755. https://doi.org/10.1016/j.immuni.2015.03.004

Sievers, F., Wilm, A., Dineen, D., Gibson, T.J., Karplus, K., Li, W., Lopez, R., McWilliam, H., Remmert, M., Söding, J., Thompson, J.D., Higgins, D.G., 2011. Fast, scalable generation of high-quality protein multiple sequence alignments using Clustal Omega. Mol. Syst. Biol. 7. https://doi.org/10.1038/msb.2011.75

Stefan, K.L., Kim, M.V., Iwasaki, A., Kasper, D.L., 2020. Commensal Microbiota Modulation of Natural Resistance to Virus Infection. Cell 183, 1312–1324.e10. https://doi.org/10.1016/j.cell.2020.10.047

Targan, S.R., Hanauer, S.B., van Deventer, S.J.H., Mayer, L., Present, D.H., Braakman, T., DeWoody, K.L., Schaible, T.F., Rutgeerts, P.J., 1997. A Short-Term Study of Chimeric Monoclonal Antibody cA2 to Tumor Necrosis Factor α for Crohn’s Disease. N. Engl. J. Med. 337, 1029–1036. https://doi.org/10.1056/nejm199710093371502

Tom, J.K., Albin, T.J., Manna, S., Moser, B.A., Steinhardt, R.C., Esser-Kahn, A.P., 2019. Applications of Immunomodulatory Immune Synergies to Adjuvant Discovery and Vaccine Development. Trends Biotechnol. https://doi.org/10.1016/j.tibtech.2018.10.004

van Nood, E., Vrieze, A., Nieuwdorp, M., Fuentes, S., Zoetendal, E.G., de Vos, W.M., Visser, C.E., Kuijper, E.J., Bartelsman, J.F.W.M., Tijssen, J.G.P., Speelman, P., Dijkgraaf, M.G.W., Keller, J.J., 2013. Duodenal Infusion of Donor Feces for Recurrent Clostridium difficile. N. Engl. J. Med. 368, 407–415. https://doi.org/10.1056/nejmoa1205037

Vatanen, T., Kostic, A.D., D’Hennezel, E., Siljander, H., Franzosa, E.A., Yassour, M., Kolde, R., Vlamakis, H., Arthur, T.D., Hämäläinen, A.M., Peet, A., Tillmann, V., Uibo, R., Mokurov, S., Dorshakova, N., Ilonen, J., Virtanen, S.M., Szabo, S.J., Porter, J.A., Lähdesmäki, H., Huttenhower, C., Gevers, D., Cullen, T.W., Knip, M., Xavier, R.J., 2016. Variation in Microbiome LPS Immunogenicity Contributes to Autoimmunity in Humans. Cell 165, 842–853. https://doi.org/10.1016/j.cell.2016.04.007

Viladomiu, M., Kivolowitz, C., Abdulhamid, A., Dogan, B., Victorio, D., Castellanos, J.G., Woo, V., Teng, F., Tran, N.L., Sczesnak, A., Chai, C., Kim, M., Diehl, G.E., Ajami, N.J., Petrosino, J.F., Zhou, X.K., Schwartzman, S., Mandl, L.A., Abramowitz, M., Jacob, V., Bosworth, B., Steinlauf, A., Scherl, E.J., Wu, H.J.J., Simpson, K.W., Longman, R.S., 2017. IgA-coated E. Coli enriched in Crohn’s disease spondyloarthritis promote TH17-dependent inflammation. Sci. Transl. Med. https://doi.org/10.1126/scitranslmed.aaf9655

Visintin, A., Mazzoni, A., Spitzer, J.H., Wyllie, D.H., Dower, S.K., Segal, D.M., 2001. Regulation of Toll-Like Receptors in Human Monocytes and Dendritic Cells. J. Immunol. 166, 249–255. https://doi.org/10.4049/jimmunol.166.1.249

Wang, Y.-H., Gorvel, J.-P., Chu, Y.-T., Wu, J.-J., Lei, H.-Y., 2010. Helicobacter pylori Impairs Murine Dendritic Cell Responses to Infection. PLoS One 5, e10844. https://doi.org/10.1371/JOURNAL.PONE.0010844

Watkins, H.C., Rappazzo, C.G., Higgins, J.S., Sun, X., Brock, N., Chau, A., Misra, A., Cannizzo, J.P.B., King, M.R., Maines, T.R., Leifer, C.A., Whittaker, G.R., DeLisa, M.P., Putnam, D., 2017. Safe Recombinant Outer Membrane Vesicles that Display M2e Elicit Heterologous Influenza Protection. Mol. Ther. 25, 989–1002. https://doi.org/10.1016/J.YMTHE.2017.01.010

Wooten, R., Ma, Y., Yoder, R., Brown, J., Weis, JH, Zachary, J., Kirschning, C., Weis, JJ, 2002. Toll-like receptor 2 is required for innate, but not acquired, host defense to Borrelia burgdorferi. J. Immunol. 168, 348–355. https://doi.org/10.4049/JIMMUNOL.168.1.348

Yang, C., Mogno, I., Contijoch, E.J., Borgerding, J.N., Aggarwala, V., Li, Z., Siu, S., Grasset, E.K., Helmus, D.S., Dubinsky, M.C., Mehandru, S., Cerutti, A., Faith, J.J., 2020. Fecal IgA Levels Are Determined by Strain-Level Differences in Bacteroides ovatus and Are Modifiable by Gut Microbiota Manipulation. Cell Host Microbe 27, 467–475.e6. https://doi.org/10.1016/j.chom.2020.01.016

Yen, D., Cheung, J., Scheerens, H., Poulet, F., McClanahan, T., Mckenzie, B., Kleinschek, M.A., Owyang, A., Mattson, J., Blumenschein, W., Murphy, E., Sathe, M., Cua, D.J., Kastelein, R.A., Rennick, D., 2006. IL-23 is essential for T cell-mediated colitis and promotes inflammation via IL-17 and IL-6. J. Clin. Invest. 116, 1310–1316. https://doi.org/10.1172/JCI21404

Zhao, S., Dai, C.L., Evans, E.D., Lu, Z., Alm, E.J., 2020. Tracking strains predicts personal microbiomes and reveals recent adaptive evolution. bioRxiv 2020.09.14.296970. https://doi.org/10.1101/2020.09.14.296970

